# The small acid-soluble proteins of *Clostridioides difficile* are important for UV resistance and serve as a check point for sporulation

**DOI:** 10.1101/2021.03.29.437554

**Authors:** Hailee N. Nerber, Joseph A. Sorg

## Abstract

*Clostridioides difficile* is a nosocomial pathogen which causes severe diarrhea and colonic inflammation. *C. difficile* causes disease in susceptible patients when endospores germinate into the toxin-producing vegetative form. The action of these toxins results in diarrhea and the spread of spores into the hospital and healthcare environments. Thus, the destruction of spores is imperative to prevent disease transmission between patients. However, spores are resilient and survive extreme temperatures, chemical exposure, and UV treatment. This makes their elimination from the environment difficult and perpetuates their spread between patients. In the model spore-forming organism, *Bacillus subtilis,* the small acid-soluble proteins (SASPs) contribute to these resistances. The SASPs are a family of small proteins found in all endospore-forming organisms, *C. difficile* included. Although these proteins have high sequence similarity between organisms, the role(s) of the proteins differ. Here, we investigated the role of the main α/β SASPs, SspA and SspB, and two annotated SASPs, CDR20291_1130 and CDR20291_3080, in protecting *C. difficile* spores from environmental insults. We found that SspA is necessary for conferring spore UV resistance, SspB minorly contributes, and the annotated SASPs do not contribute to UV resistance. In addition, none of these SASPs contribute to the resistance of tested chemicals. Surprisingly, the combined deletion of *sspA* and *sspB* prevented spore formation. Overall, our data indicate that UV resistance of *C. difficile* spores is dependent on SspA and that SspA and SspB regulate / serve as a checkpoint for spore formation, a previously unreported function of SASPs.

**Importance:** *C. difficile* infections remain problematic and elimination of spores within an environment is essential to limit person-to-person spread. A deeper understanding of how spores resist cleaning efforts could lead to better strategies to eradicate the spores in a contaminated environment. The small acid-soluble proteins (SASPs), found in all endospore-forming organisms, are one mechanism that allows for spore resilience. Here, we find that *C. difficile* SspA and SspB protect against UV light. Unexpectedly, these SASPs also regulate spore formation, a role not described for any SASP to date.

## Introduction

*Clostridioides difficile* is the leading cause of antibiotic associated diarrhea with ∼224,000 annual infections in the United States [1–3]. Prior antibiotic treatment is the greatest risk factor for *C. difficile* infection due to their broad-spectrum activity that can lead to a dysbiotic colonic microbial community [4, 5]. Upon inoculation into a susceptible host, *C. difficile* spores germinate to the toxin-producing vegetative form [4]. These toxins lead to the disruption of the colonic epithelium and the common symptoms of disease (*e.g.*, diarrhea and colitis) [4]. Vancomycin and fidaxomicin are the recommended treatments for *C. difficile* infection [4, 6]. While these antibiotics treat the infection by targeting the actively-growing vegetative cells, the spore form is resistant to antibiotics [5, 7]. In contrast to the anaerobic nature of the *C. difficile* vegetative cell, spores can survive the oxygen-rich environment outside of a host, and are considered the transmissible form [5, 8–10].

Endospores are dormant forms of bacteria that can withstand extreme environmental conditions and chemical exposures [11]. In all endospore-forming bacteria, sporulation is controlled by the master transcriptional regulator, Spo0A [10]. Activation of Spo0A by phosphorylation results in the global activation of gene expression with the end goal of optimizing / surviving post-exponential phase growth [12–14]. Upon the initiation of sporulation, the vegetative cell divides asymmetrically, resulting in a large mother cell compartment and a smaller forespore compartment [13]. Subsequently, a cascade of sigma factor activation occurs and leads to the development of a dormant endospore [13]. In *Bacillus subtilis*, a model organism for studying sporulation and germination, the activation of σ^F^ in the forespore, leads to σ^E^ activation in the mother cell. The activation of σ^E^ in the mother cell, in turn, leads to activation of σ^G^ in the forespore and then results in the activation of σ^K^ in the mother cell. The result of this crisscross sigma factor activation cascade is the engulfment of the forespore by the mother cell, maturation of the forespore, and the eventual release of the spore by lysis of the mother cell; the same sigma factors drive *C. difficile* sporulation but the crisscross activation across compartments does not occur [12, 13, 15–17]. The resulting spores are extremely resistant to environmental conditions and common cleaning methods [11, 18]. Thus, with a deeper understanding of the resistance properties of spores, and the mechanisms that confer this resistance, novel interventions could be developed to clean contaminated environments.

The small acid-soluble proteins (SASPs) confer resistance to spores [11, 19]. The SASPs are a family of proteins that are less than 100 amino acids in length and are conserved among all endospore forming organisms [19]. They are produced late in sporulation under the forespore-specific sigma factor, σ^G^, and account for approximately 20% of the total spore protein content [11, 16, 19]. Most spore-forming bacteria encode the two major α/β-type SASPs (SspA and SspB), however there are other minor SASPs that vary in number depending on the organism [19–23]. *B. subtilis* also encodes a γ-type SASP that is hypothesized to serve as an amino acid reservoir for use upon outgrowth of the vegetative cell from the germinated spore [24]. Clostridial species, to date, have not been found to contain γ-type SASPs [20, 21]. In *C. difficile,* the R20291 strain encodes *sspA* and *sspB* orthologues and two genes annotated as putative SASPs, *CDR20291_1130* and *CDR20291_3080*. SASPs are well-conserved across genera with 70% similarity between *C. difficile* and *B. subtilis* SspA and SspB [25]. However, the annotated SASPs have less similarity to *C. difficile* or *B. subtilis* SspA and SspB, ranging from 40-60%. Interestingly, CDR20291_3080 shares 80% similarity to *C. perfringens* Ssp4, a novel SASP, which protects the spores from nitrous acid and extreme heat associated with food processing [23, 26].

In *B. subtilis*, the α/β-type SASPs contribute to spore resistance against several chemicals, such as nitrous acid, formaldehyde, glutaraldehyde, iodine, or hydrogen peroxide [18, 26–34]. Moreover, *B. subtilis* Δ*sspA* mutants completely lose viability after 3 minutes of exposure to 254 nm UV light, and *B. subtilis* Δ*sspB* mutants have 10% survival after 7 minutes of exposure [35]. Interestingly, a double mutant was even more sensitive to UV exposure than were vegetative cells, highlighting the importance of these proteins in spore survival [35]. These UV-sensitive phenotypes could be complemented by expressing, *in trans*, either *sspA* or *sspB* [36]. Moreover, a SASP from *B. megaterium* complemented the phenotype, suggesting that they could play interchangeable roles in UV resistance [31, 36].

In other spore-forming bacteria, the role of the SASPs vary among organisms. In *C. botulinum,* SASPs were found to be necessary for protection against nitrous acid, similar to what is observed in *B. subtilis* [28]. However, they are not necessary for protection against hydrogen peroxide or formaldehyde, contrary to what is observed in *B. subtilis* [28]. In *C. perfringens* isolates that cause food poisoning, Ssp4 was necessary for spores surviving food processing events (*e.g.*, high heat and use of nitrites) [23, 26]. Other *C. perfringens* SASPs were found to protect spores against UV light, hydrogen peroxide, nitrous acid, formaldehyde, and hydrochloric acid [29, 37, 38].

The α/β-type SASPs of *B. subtilis, C. perfringens,* and *C. acetobutylicum* all bind to DNA *in vitro* [37, 39–41]. In *B. subtilis*, the binding of these proteins changes the confirmation of the DNA to one between an A and a B form [39, 41–43]. In this unique conformation, an alternative form of UV damage is induced, a thymidyl-thymidine adduct, called the spore photoproduct. The spore photoproduct is repaired by the spore photoproduct lyase, SPL, during germination and outgrowth [25, 44–48].

Here, we investigate the functions of *C. difficile sspA*, *sspB*, and the annotated SASPs, *CDR20291_1130* and *CDR20291_3080.* We found that *C. difficile* SspA is the major contributor to UV resistance of the spores and that SspB is minorly involved in UV resistance. CDR20291_1130 and CDR20291_3080 do not contribute to UV resistance. Additionally, none of the SASPs contribute to spore chemical resistance in the concentrations and exposure times tested. Surprisingly, we found that the deletion of both *sspA* and *sspB* prevented spore formation. Our results indicate that, in addition to providing UV resistance to spores, the major *C. difficile* SASPs are involved in spore formation.

## Results

### Conservation of SASPs in C. difficile

The main α/β type SASPs are conserved in spore-forming bacteria [19]. Within *C. difficile*, SspA and SspB are 85% identical. The putative SASPs are approximately 40% identical to the main α/β type SASPs. All 4 contain the conserved “EIA” sequence that is cleaved by the germination protease (GPR) upon germination (Figure 1) [49–51]. Even with strong sequence similarity / identity, SASPs have varying roles between spore forming organisms. To investigate the roles of *C. difficile sspA*, *sspB*, *CDR20291_1130*, and *CDR20291_3080* in UV resistance, outgrowth, and chemical resistance, single deletions and pairwise deletions were generated using CRISPR-Cas9 genome editing [52]. Total amounts of spores produced and the sporulation rates by the single deletion strains, *C. difficile* Δ*sspA, C. difficile* Δ*CDR20291_1130,* and *C. difficile* Δ*CDR20291_3080* were indistinguishable from the wildtype *C. difficile* R20291 parental strain (Figure S1). Surprisingly, the *C. difficile* Δ*sspB* strain did not make spores and will be discussed later. These results indicate that the single deletions of *C. difficile sspA, C. difficile CDR20291_1130* and *C. difficile CDR20291_3080* do not impact *C. difficile* sporulation.

**Figure 1.**
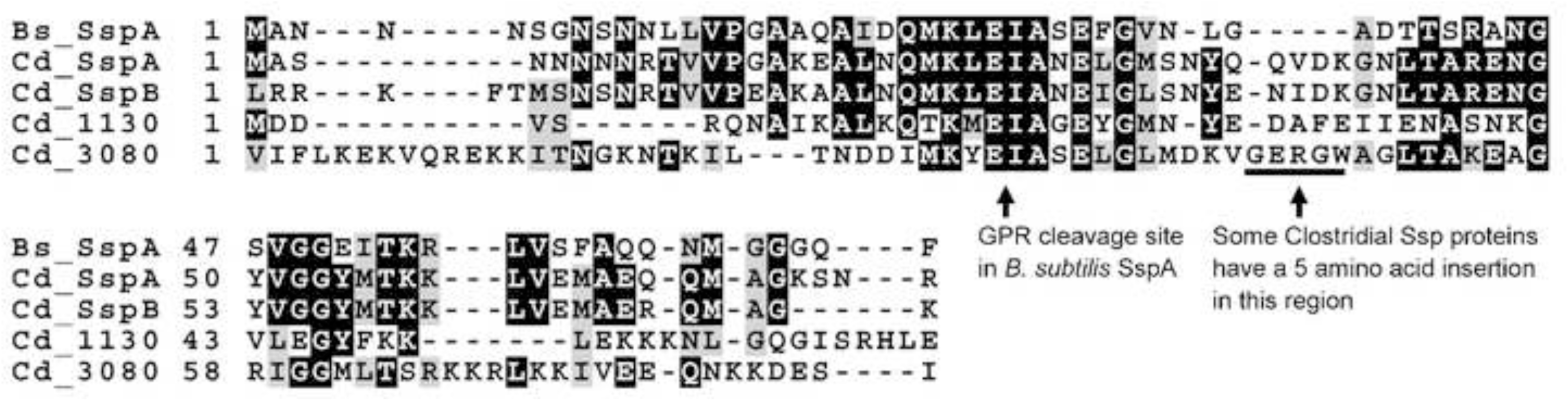
SASP homology. Alignment of *B. subtilis* SspA with *C. difficile* SspA, SspB, CDR20291_1130 and CDR20291_3080. The site of GPR cleavage is indicated. Also illustrated is the site of insertion in Clostridial Ssp proteins.

### sspA is required for spore UV resistance

Because of the strong phenotype of *B. subtilis* SASPs in UV resistance, we hypothesized that *C. difficile* SspA, and / or the two annotated SASPs (CDR20291_1130 and CDR20291_3080) may function similarly. The viability of SASP mutants after a 10 minute exposure to 302 nm UV light was tested. Viable spores were quantified by colony formation on media supplemented with taurocholate (a *C. difficile* spore germinant) and then compared to spores derived from the wildtype parental strain [53]. Exposure of the *C. difficile* Δ*sspA* strain to UV light resulted in a ∼1,000x decrease in spore survival. Spores derived from the *C. difficile* Δ*CDR20291_1130* and *C. difficile* Δ*CDR20291_3080* mutants had a statistically significant difference in UV resistance compared to spores derived from the wildtype strain, but this difference is likely not biologically relevant (Figure 2A). These results indicate that *C. difficile* SspA is the most important SASP for UV resistance while the genes annotated as putative SASPs are not involved in UV resistance. In *B. subtilis*, SASP binding to DNA helps to protect from UV by encouraging the formation of spore photoproducts (SP) instead of cyclobutane thymine dimers [18, 54]. The spore photoproduct lyase, SPL, repairs the SP [44–46, 55]. To understand the impact of SPL on UV resistance in *C. difficile,* we engineered a deletion in the *C. difficile spl* gene. Spores derived from *C. difficile* Δ*spl* strain have approximately 10x reduced survival compared to spores derived from the wildtype *C. difficile* R20291 strain (Figure 2A). Unsurprisingly, the *C. difficile* Δ*sspA* Δ*spl* double mutant remains at the level of *sspA* mutant alone (0.1% of wildtype), suggesting that SPL functions in a capacity where without SspA, SPL is not necessary.

**Figure 2.**
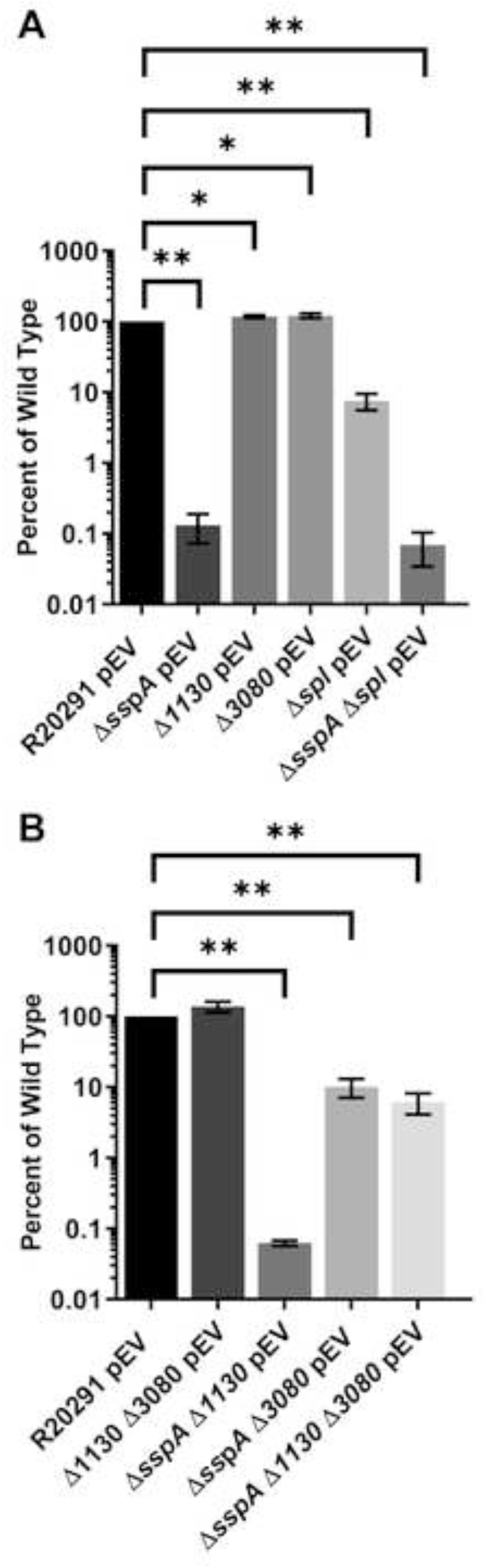
*C. difficile sspA* is necessary for UV protection. Spores were exposed to 302 nm UV light for 10 minutes. After exposure, they were serially diluted and plated onto rich medium supplemented with germinants and CFUs were enumerated. All data was normalized to T_0_ then the ratio of the mutants was normalized to the ratio of wildtype. A.) Spores derived from wild type R20291 with the indicated deletion strains. B.) spores derived from pairwise deletions of *C. difficile* Δ*sspA*, Δ*CDR20291_1130*, and Δ*CDR20291_3080.* pEV indicates an empty vector. All data points represent the average from three independent experiments. Statistical analysis was performed by one way ANOVA with Dunnett’s multiple comparisons test in comparison to wildtype. * P< 0.01, ** P< 0.0001

To determine if the putative SASPs have redundant roles in UV resistance, we generated deletions of all pairwise combinations of *C. difficile CDR20291_1130, C. difficile CDR20291_3080,* and *C. difficile sspA*. After 10 minutes of UV exposure, spores derived from the *C. difficile* Δ*CDR20291_1130* Δ*CDR20291_3080* were as resistant as wildtype spores, indicating that these annotated SASPs are not compensating for each other during UV exposure (Figure 2B). Spores derived from the *C. difficile* Δ*sspA* Δ*CDR20291_1130* mutant had no further reduction in survival compared to the *C. difficile* Δ*sspA* mutant alone. However, spores derived from the *C. difficile* Δ*sspA* Δ*CDR20291_3080* double mutant and the Δ*sspA* Δ*CDR20291_1130* Δ*CDR20291_3080* triple mutant were not as sensitive to UV light as the *C. difficile* Δ*sspA* or the *C. difficile* Δ*sspA* Δ*CDR20291_1130* strains (Figure 2B).

### The sspA promoter is necessary for complementation of C. difficile ΔsspA UV resistance

To understand the extent of protection that *C. difficile* SspA provides against UV damage, we quantified the viability of spores exposed to UV over time. Spores derived from the *C. difficile* Δ*sspA* strain showed a 10% loss of viability after 2.5 minutes of UV exposure. This loss increased to approximately 3 log_10_ after 15 minutes of exposure. Expression of *sspA* from a plasmid under the control of its native promoter restored viability to the *C. difficile* Δ*sspA* strain (Figure 3A).

**Figure 3.**
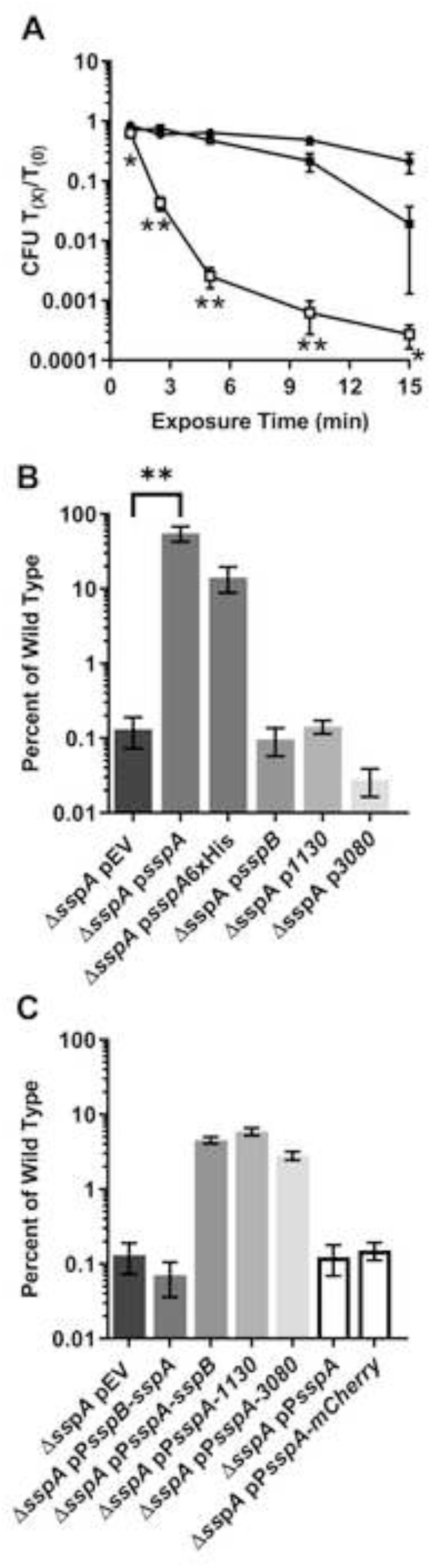
Complementation of the *C. difficile* Δ*sspA* strain. A) Spores derived from wildtype *C. difficile* R20291 pEV ●, the *C. difficile* Δ*sspA* pEV deletion strain □, and the *C. difficile* Δ*sspA* p*sspA* strain ▪ were exposed to UV for varying times and the viability assessed by normalizing to T_0_. B) Spores from the *C. difficile* Δ*sspA* mutant strain containing genes under control of the *sspA* promoter were exposed to 302 nm UV light for 10 minutes. After exposure, the spores were serially diluted and plated onto rich medium supplemented with germinants. The CFUs were enumerated and then normalized to T_0_ CFUs. The ratio of the complemented *C. difficile* Δ*sspA* mutant strains were normalized to the ratio of the wildtype strain. C) Spores from the *C. difficile* Δ*sspA* mutant strain containing genes under the control of the *sspA* or *sspB* promoter regions were exposed to UV for 10 minutes. pEV indicates an empty vector. All data represents the average of three independent experiments. Statistical analysis was performed by ANOVA with Dunnett’s multiple comparison. A) two way, in comparison to wildtype, B) and C) one way, in comparison to *C. difficile* Δ*sspA* pEV. * P<0.05, ** P<0.0001.

To further understand the role of *C. difficile* SspA in UV resistance, we tested the impact of different *sspA* expression constructs on spore survival. The *sspA* complement consisting of the *sspA* gene under its native promoter region resulted in restoration of spore viability. A 6x-histidine tag inserted on the C-terminus of *sspA* also resulted in restoration of spore viability upon UV exposure (Figure 3B). In *B. subtilis,* SASP genes can cross-complement a SASP mutant [36]. Therefore, plasmids were constructed that consisted of *C. difficile sspB, CDR20291_1130,* or the *CDR20291_3080* genes, driven by their native promoter regions, introduced into the *C. difficile* Δ*sspA* mutant strain. After 10 minutes of exposure to UV light, spores derived with these plasmid constructs revealed that *C. difficile sspB, CDR20291_1130,* or *CDR20291_3080* were unable to restore the UV resistance to spores derived from the Δ*sspA* mutant strain (Figure 3B). To determine if this is an issue with differences in expression, the promoter regions were changed and these genes were again tested for their ability to complement the *C. difficile* Δ*sspA* strain. When the *sspA* gene was placed under control of the *sspB* promoter region, complementation no longer occurred. This further supported our hypothesis of the SASPs containing differences in expression levels. Swapping the promoter regions of *C. difficile sspB, CDR20291_1130*, or *CDR20291_3080* complementation plasmids for the *C. difficile sspA* promoter region resulted in a restoration to approximately 5% of wildtype levels (Figure 3C). However, as negative controls, the *sspA* promoter region alone, or the *sspA* promoter driving the gene encoding mCherry, could not complement the UV phenotype. These results suggest that despite what is observed for cross-complementation in other organisms, *C. difficile sspB*, *CDR20291_1130*, or *CDR20291_3080* cannot fully complement the *C. difficile* Δ*sspA* strain phenotype, and to provide any complementation, they must be expressed from the *sspA* promoter.

### C. difficile SASPs have redundant functions during outgrowth

*C. difficile CDR20291_1130* and *CDR20291_3080* do not contribute to spore UV resistance, but it is possible that their main role is to protect against other harsh environmental conditions or to serve as amino acid reservoirs (like γ-type SASPs of *B. subtilis*) during outgrowth of a vegetative cell from the germinated spore [24]. To determine if these annotated SASPs contribute to outgrowth, the OD_600_ of spores derived from wildtype and mutant strains were analyzed over 12 hours in complex medium supplemented with germinants [53, 56]. No difference in the outgrowth of vegetative cells was observed for the mutant strains, compared to the wildtype parental strain.

It is possible that the complex medium masked the hypothesized phenotype due to the sheer abundance of nutrients in the medium eliminating the need for the amino acids derived from the SASPs. Instead, we tested if outgrowth of a spore in minimal medium would be influenced in these mutant strains. Unfortunately, using a minimal medium resulted in an extreme delay in outgrowth that was not possible to practically measure. Therefore, spore outgrowth was analyzed in half-strength complex medium. In addition, to eliminate the possibility of extra resources being packaged into the spore when grown on a complex medium, and thus reducing the need for SASPs during outgrowth, spores were generated on minimal medium. Again, despite spore production on minimal medium and using half-strength complex medium during the assay, there were no differences between the outgrowth of wildtype spores or spores derived from the *C. difficile* Δ*sspA* or the *C. difficile* Δ*CDR20291_1130* Δ*CDR20291_3080* double mutant strain (Figure 4A).

**Figure 4.**
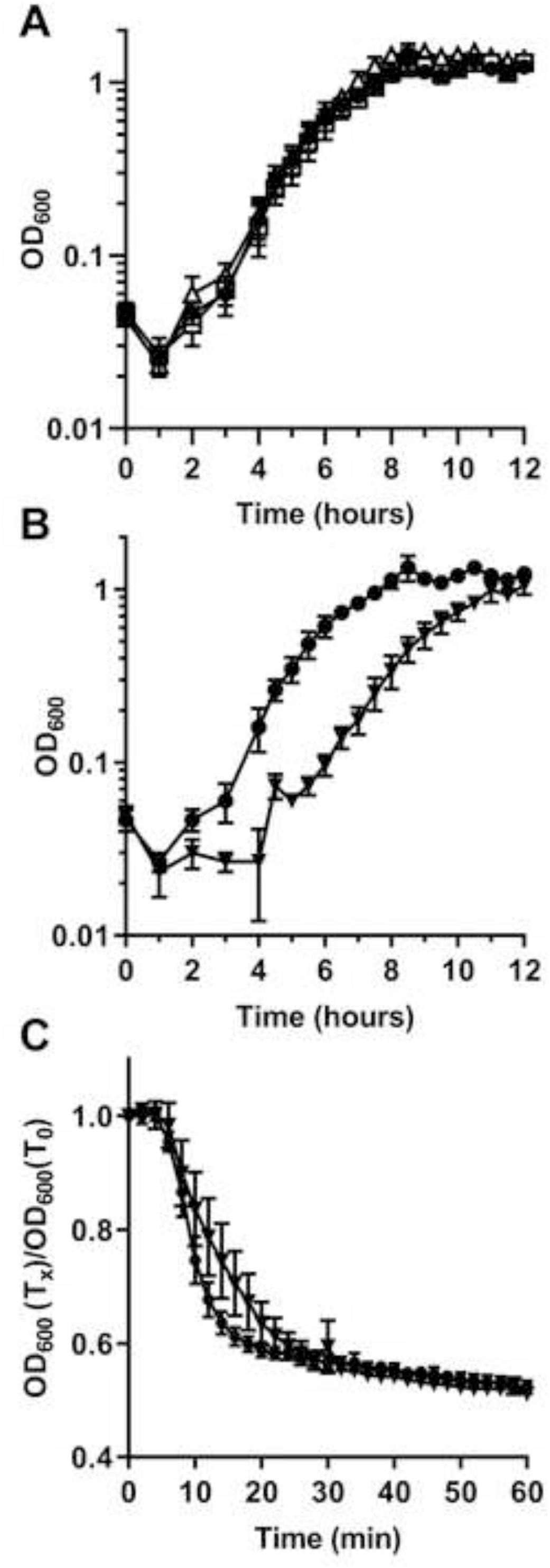
*C. difficile* Δ*sspA* Δ*CDR20291_1130* Δ*CDR20291_3080* affects spore outgrowth. Outgrowth was determined by monitoring the OD_600_ of spores inoculated into half-strength complex medium supplemented with germinants (10 mM taurocholic acid and 30 mM glycine). All spores used for outgrowth were generated on minimal medium. A) Outgrowth of of *C. difficile* R20291 pEV ●, *C. difficile* Δ*sspA* pEV □ and *C. difficile* Δ*CDR20291_1130* Δ*CDR20291_3080* pEV Δ. B) Outgrowth of wildtype R20291 pEV ● and triple mutant spores of *C. difficile* Δ*sspA* Δ*CDR20291_1130* Δ*CDR20291_3080* pEV ▾. C) Spore germination was monitored at OD_600_ upon exposure of spores to germinants taurocholic acid and glycine in a buffered solution. The OD_600_ was normalized to T_0_. R20291 pEV ● and triple mutant spores of *C. difficile* Δ*sspA* Δ*CDR20291_1130* Δ*CDR20291_3080* pEV ▾. pEV indicates an empty vector. All data points represent the average from three independent experiments. Statistical analysis by two way ANOVA with Dunnett’s multiple comparisons test. A) *C. difficile* Δ*sspA,* P<0.001 at 11 minutes.

Due to the possibility that the SASPs can compensate for the deletion of one, a triple mutant was generated of *sspA, CDR20291_1130,* and *CDR20291_3080.* These spores were also generated on minimal medium and outgrowth analyzed in half-strength complex media. Spores derived from the *C. difficile* Δ*sspA* Δ*CDR20291_1130* Δ*CDR20291_3080* triple mutant had an approximate 2-hour delay in outgrowth compared to the wildtype strain (Figure 4B). To eliminate the possibility that the difference in outgrowth is due to a germination defect in this triple mutant, we monitored germination by OD_600_ in buffer supplemented with germinants [53, 56]. During the very early events of endospore germination, the dormant, phase bright, spore transitions to a phase dark, germinated spore. Mutant spores germinated similarly to wildtype, suggesting no defect in germination (Figure 4C). These results suggest that *C. difficile* SspA, CDR20291_1130, and CDR20291_3080 could be used as a nutrient / amino acid source during outgrowth of a germinated spore.

### The C. difficile SASPs do not contribute to chemical resistance

To further characterize the role of these proteins in the spore, spores were exposed to various chemicals. Spores derived from the single deletion of *C. difficile sspA*, *CDR20291_1130*, and *CDR20291_3080* and their complements, when necessary, were exposed to chemicals for 1 minute, 5 minutes, 10 minutes, and 30 minutes and spore viability was assessed by plating onto rich medium supplemented with germinant. Colony forming units were compared to T_0_ and then this ratio compared to the ratio of wildtype survival. Spores exposed to 3% H_2_0_2_, 75% EtOH, 0.25% glutaraldehyde, 1 M HCl, 0.05% hypochlorite, and 2.5% formaldehyde did not exhibit reduced viability in comparison to spores derived from the wildtype strain (Figure S2A-F). The mutant strains did have a slight reduction in viability after 5 or 10 minutes of exposure to 250 mM nitrous acid (Figure S2G).

### C. difficile spore formation is influenced by SASPs

When generating the *C. difficile* Δ*sspB* mutant strain, we found that spores derived from this strain were phase gray and were not released from the mother cell, compared to the wildtype strain (Figure 5A and 5B). This phenotype could be complemented by expression of the *sspB* gene under the control of the *sspB* promoter region (Figure 5C) or the *sspA* gene under control of the *sspA* promoter region (Figure 5D). However, only a σ^G^-controlled promoter was able to complement the phenotype, other σ-factor controlled promoters could not complement (Figures 5E - H). The rate of sporulation of the *C. difficile* Δ*sspB* mutant was also 5 log_10_ less than the wildtype strain and was complemented by expression of *sspB* from a plasmid (Figure S3). Surprisingly, whole genome resequencing of the *C. difficile* Δ*sspB* strain revealed an additional, single nucleotide mutation in *sspA*. This mutation resulted in an *sspA*_G52V_ allele, in addition to the *sspB* deletion (referred to as *sspB*_SNP_).

**Figure 5.**
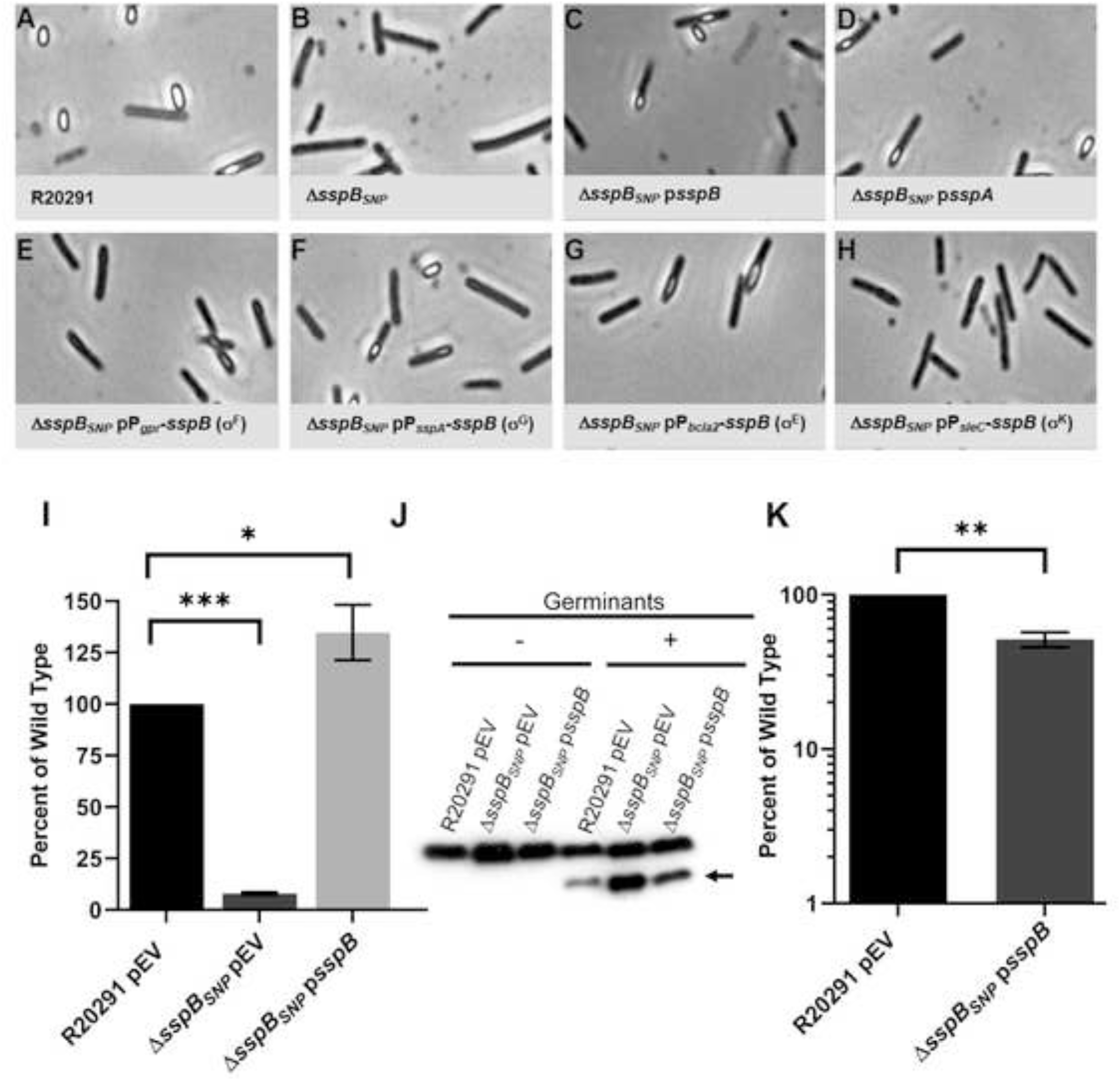
*C. difficile* Δ*sspB_SNP_* results in phase gray engulfed spores. A-H) Day 6 cultures of the indicated strains were fixed in 4% formaldehyde and 2% glutaraldehyde before imaging on a Leice DM6B microscope. I) DPA content was determined by boiling spore solutions and analyzing approximately 2.5 x 10^5^ spores by terbium florescence. J) Western blot of spores exposed to 10 mM taurocholic acid in a rich medium for 1 hour and blotted against SleC. ← indicates processed SleC. K) Spores were exposed to 302 nm UV light for 10 minutes before serial dilution and plating on rich medium supplemented with germination. All values were normalized to T_0_ and then those ratios were normalized to the ratio of wildtype survival. pEV indicates an empty vector. All data points represent the average from three independent experiments. I) Statistical analysis by one way ANOVA with Dunnetts multiple comparisons test. K) Statistical analysis by unpaired t test. * P<0.05, ** P<0.01, *** P<0.001.

Due to the phenotype of this strain, normal spore purification processes were unsuccessful. To encourage release of the immature spores from the mother cells, cultures from sporulating cells were incubated with lysozyme before resuming normal spore purification steps. This encouraged the spores to release from the mother cells and the recovery of a limited number of phase gray spores. The phase bright phenotype of wildtype spores is partially attributed to the dipicolinic acid (DPA) content of the spore core. Approximately 2.5 x 10^5^ spores of the *C. difficile* wildtype, Δ*sspB*_SNP_, and Δ*sspB*_SNP_ p*sspB* strains were boiled and the DPA levels determined by Tb^3+^ fluorescence [57]. The DPA content of the *C. difficile* Δ*sspB*_SNP_ strain was significantly lower than wildtype content and this was able to be complemented by expression of the *sspB* gene from its native promoter region (Figure 5I).

Because germination assays rely heavily on the release of DPA during germination (phase bright to phase dark transition), we were unable to use these assays to determine if this deletion and SNP combination altered germination capabilities [57, 58]. Instead, the spores were exposed to germinants and the processing of spore proteins during germination were analyzed by immunoblot. In response to germinants, spores derived from the *C. difficile* Δ*sspB*_SNP_ strain processed proSleC to its active form, indicating that they are capable of receiving the germinant signals (Figure 5J) [59–61].

Finally, we tested the ability of spores derived from the *C. difficile* Δ*sspB*_SNP_ p*sspB* strain to resist UV damage. The *C. difficile* Δ*sspB*_SNP_ alone was not assessed due to the difficulty in producing and purifying spores. In the assay, the *C. difficile* Δ*sspB*_SNP_ p*sspB* strain was complemented to wildtype levels, revealing that the *sspA*_G52V_ allele is not impaired in its ability to protect against UV damage but SspA may regulate *C. difficile* sporulation with SspB (Figure 5K).

### C. difficile sspA and sspB are required for spore formation

Due to the G52V mutation in *sspA*, we created a clean *C. difficile* Δ*sspA* Δ*sspB* strain to determine if the double mutant had the same phenotype as the *C. difficile* Δ*sspB*_SNP_ strain. Similar to above, the *C. difficile* Δ*sspA* Δ*sspB* strain produced phase gray, immature spores (Figure 6A). The rate of sporulation was also 1,000x less than wildtype and could be complemented with expression of *sspA* and *sspB* from a plasmid (Figure S3). Moreover, analysis of the DPA content also showed little DPA in the double mutant strain, in comparison to wildtype (Figure 6B). Finally, to evaluate whether these spores can germinate, we monitored SleC activation. These double mutant spores also processed proSleC to the active SleC form, showing that they are still able to germinate (Figure 6C) [59–61]. Again, due to the difficulty in purifying this strain and despite density gradient purification, spore preparations still contained debris making it difficult to quantify the spores by microscopy. Therefore, we were forced to use approximate spore counts. The Coommassie stained SDS PAGE gel shows that more protein is present in the double mutant sample than in wildtype or the complemented strains, even though the SleC bands detected by western indicate similar concentrations (Figure S4). This is possibly due to the presence of contaminating vegetative cells in the double mutant preparation because of the difficulty in purifying or the spore counts could have been underestimated.

**Figure 6.**
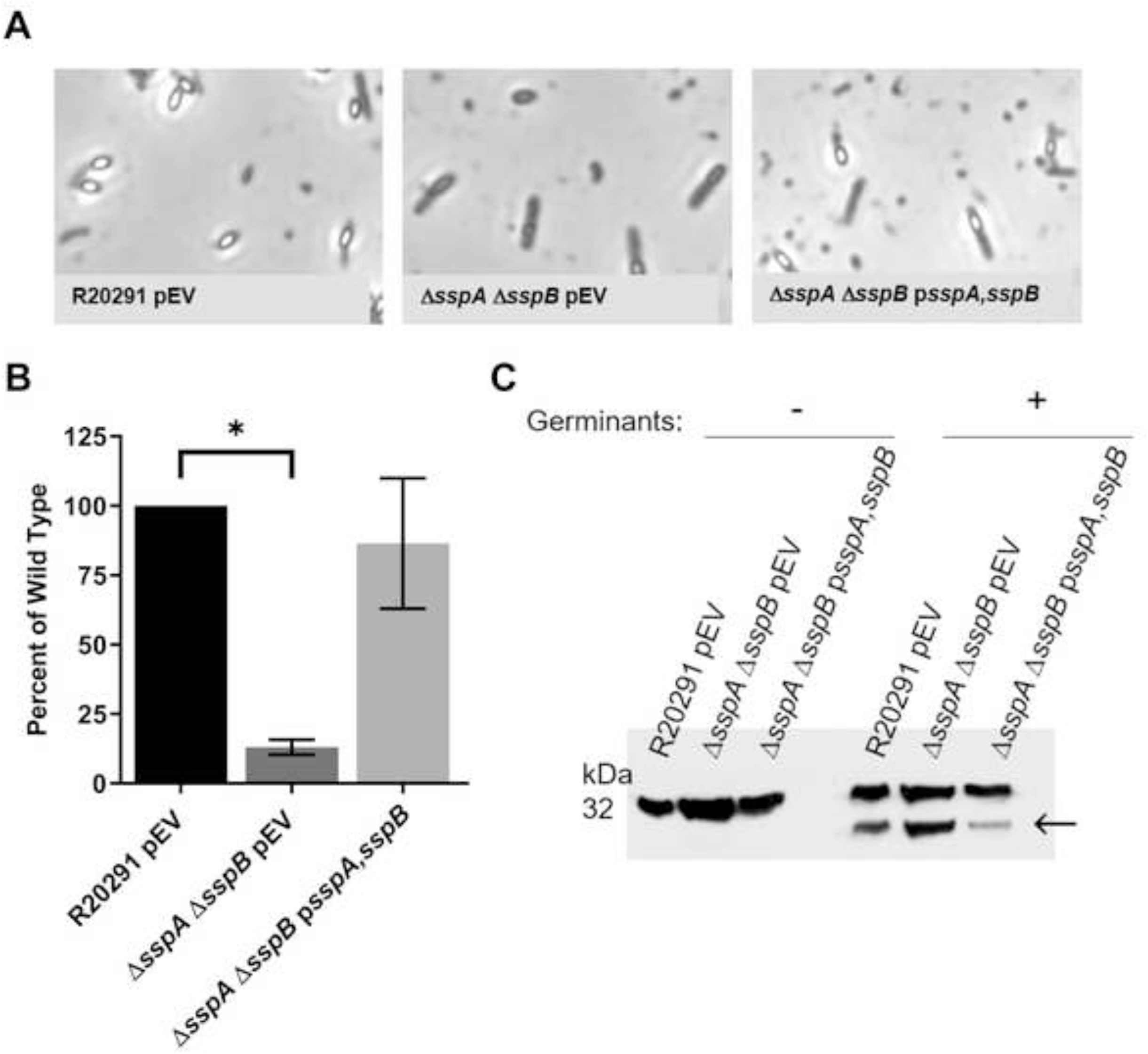
Phenotypic characterization of *C. difficile* Δ*sspA* Δ*sspB* double mutant. A) Day 6 cultures were fixed in 4% formaldehyde and 2% glutaraldehyde before imaging on a Leica DM6B microscope. B) DPA content was determined by boiling spore solutions and analyzing approximately 2.5 x 10^5^ spores by terbium florescence. C) Western blot of spores exposed to 10 mM taurocholic acid in a rich medium for 1 hour and blotted against SleC. ← indicates the cleaved SleC band. pEV indicates an empty vector. Data represents the average of three independent experiments. Statistical analysis was performed by one way ANOVA with Dunnett’s multiple comparison test. * P<0.01.

### C. difficile sspB plays a minor role in UV resistance and does not play a role in outgrowth or chemical resistance

After discovering the SNP in the *C. difficile* Δ*sspB* strain, we generated a clean deletion that produces phase bright, released spores. The *C. difficile* Δ*sspB* strain had a sporulation rate identical to wildtype (Figure S3).

Since a clean deletion produces spores, the UV resistance could be determined. Spores derived from the wildtype *C. difficile* R20291, *C. difficile* Δ*sspB* mutant, and *C. difficile* Δ*sspB* p*sspB* strains were exposed to UV light for 10 minutes before plating on media with germinants to determine CFUs. Spores derived from the *C. difficile* Δ*sspB* strain had approximately 10% of the resistance observed for the wildtype strain. This phenotype was complemented by expression of *sspB* from its native promoter *in trans* (Figure 7A). Therefore, *C. difficile* SspB plays a minor role in UV resistance.

**Figure 7.**
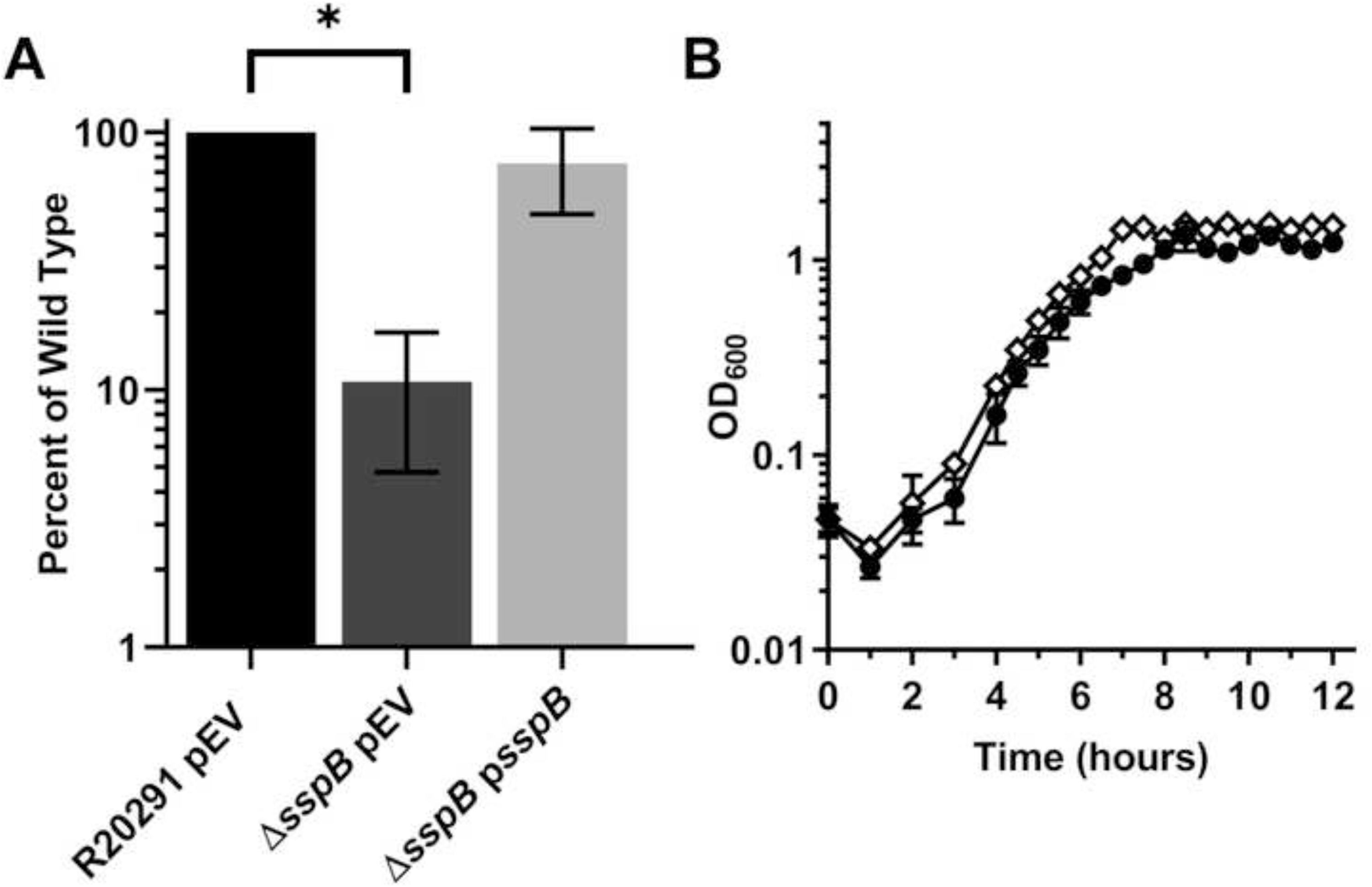
*C. difficile* Δ*sspB* has a minor UV defect and no role in outgrowth. A) Spores derived from *C. difficile* R20291 pEV, *C. difficile* Δ*sspB* with pEV, and *C. difficile* Δ*sspB* p*sspB* were exposed to UV light for 10 minutes. The spores were serially diluted and plated onto medium containing germinants. Strains were normalized to CFUs at T_0_. The ratios were then normalized to the ratio of wildtype survival. B) Outgrowth of spores from *C. difficile* R20291 pEV and *C. difficile* Δ*sspB* pEV **◊** was monitored at OD_600_ over 12 hours. Starting OD was 0.05 into half-strengthed complex medium supplemented with germinants. pEV indicates an empty vector. All data represents the average of three independent experiments. A) Statistical analysis by one way ANOVA with Dunnett’s multiple comparison test. * P<0.001. B) Statistical analysis by two way ANOVA with Dunnett’s multiple comparison test, P<0.0001 at 10 hours.

Next, we tested the outgrowth of spores derived from the *C. difficile* Δ*sspB* strain. Again, the spores were generated on a minimal medium and the outgrowth was tested in half-strength complex media supplemented with germinants. Similar to our observations with the Δ*sspA* strain, there was no difference in the outgrowth of spores from the *C. difficile* Δ*sspB* strain in comparison to wildtype spores (Figure 7B).

Spores from *C. difficile* Δ*sspB* were also tested for chemical resistance in the same manner as previously discussed. In comparison to *C. difficile* R20291 wildtype spores, *C. difficile* Δ*sspB* spores exposed to 3% H_2_0_2_, 75% EtOH, 0.25% glutaraldehyde, 1 M HCl, 0.05% hypochlorite, 2.5% formaldehyde, and 250 mM nitrous acid did not have reduced viability (Figure S2A-G).

## Discussion

*C. difficile* infections occur in individuals with a disrupted microbiota, commonly due to the use of broad-spectrum antibiotics [1, 4, 5]. Vegetative *C. difficile* colonizes the lower GI tract and produces toxins that disrupt the epithelial lining integrity and cause colitis and diarrhea [4]. While the toxin producing vegetative cells are eliminated with antibiotic treatment, the spores can persist in the gut and become vegetative cells again, or they can shed into the environment [4, 5]. This shedding allows for the transmission of disease by passing spores into an environment where they can then transfer to other individuals [4, 9]. Spores can withstand environmental insults, such as UV light, and many chemicals, and persist long term in most environments [11, 18, 25].

UV-B light (between 315 – 280 nm) is the UV wavelength range most responsible for DNA breaks and DNA lesions (*e.g.,* cyclobutane pyrimidine dimers) [62]. DNA does not absorb UV-A (400 – 315 nm) but UV-C (less than 280 nm) has high energy and is known to cause some DNA mutations, but fewer cyclobutane pyrimidine dimers than UV-B [62]. Though *C. difficile* spores are more likely to encounter UV-B from environmental UV radiation, UV-C is absorbed by ozone making it less likely that spores encounter this spectrum in nature [62]. Several studies have shown that treatment with 254 nm UV light reduced viable *C. difficile* spore load in the hospital environment [63, 64]. However, this method is not ideal due to the potential exposure of dangerous UV-C light to individuals implementing this cleaning technique and the long exposure time of approximately 1 hour. An interesting study evaluated the effects of a hospital biocide, sodium dichloroisocyanurate (NaDCC), on *C. difficile* spores. In this study, a 10x less lethal concentration of biocide was tested [65]. At this concentration, spores are not killed as effectively as the lethal concentration and the hydrophobicity of the spores was greatly reduced (to approximately 20%) [65]. The reduction in hydrophobicity is of concern because it is possible that spores would spread easier due to reduced adherence to surfaces. This further perpetuates the problem of trying to clean a *C. difficile* contaminated environment.

The small acid-soluble proteins (SASPs) are well established to protect the spores from environmental insults (*e.g.*, UV light or chemicals) [11, 18, 19, 25]. Here we investigated the functions of *C. difficile* SASPs in response to UV light and chemicals. We found that *C. difficile* SspA and SspB have a role in UV resistance but none of the SASPs significantly contribute to chemical resistances. Surprisingly, we discovered that the deletion of *C. difficile* Δ*sspA* Δ*sspB* resulted in phase gray, unreleased spores and we hypothesize that SspA and SspB are working together to influence sporulation, a SASP function not previously reported.

*C. difficile* SASPs are involved in protecting spores from UV exposure. This was not surprising since the SASPs in all other organisms have had a large role in UV resistance [29, 35, 38, 66, 67]. While *B. subtilis* Δ*sspB* had almost 100% survival after 3 minutes of exposure to 254 nm UV, *B. subtilis* Δ*sspA* spores were sensitive to UV light with only 0.1% survival [35]. In *C. difficile*, SspA is the major contributor to UV resistance. *C. difficile* Δ*sspA* spores exposed to UV 302 nm light for 10 minutes had 0.1% survival in comparison to *C. difficile* Δ*sspB* which had 10% survival; *C. difficile* Δ*CDR20291_1130* and *C. difficile* Δ*CDR20291_3080* did not lose viability. UV protection is contributed to the binding of the SASPs to the DNA, which changes its conformation and encourages spore photoproduct mutations over cyclobutane pyrimidine dimers [19, 25, 39, 41, 44, 46, 54, 68]. The loss of the spore photoproduct repair system, SPL, interestingly only resulted in a 10x loss of viability. This highlights the importance of SspA in protecting the genome from lethal UV irradiation. However, in a *C. difficile* Δ*sspA* Δ*spl* double mutant, the viability remained the same as for a *C. difficile* Δ*sspA* single deletion, suggesting SspA is the major contributor to UV resistance. This also suggests that SspA primarily influences the change in DNA structure that encourages spore photoproduct formation upon UV exposure.

In *B. subtilis,* other SASPs than SspA and SspB are considered minor SASPs with many redundant roles [19, 25, 36]. To evaluate if the *C. difficile* annotated SASPs can compensate for each other or only minorly contribute to UV resistance, pairwise deletions were generated. The annotated SASPs do not compensate for each other, as a *C. difficile* Δ*CDR20291_1130* Δ*CDR20291_3080* double deletion does not have a UV defect. A *C. difficile* Δ*sspA* Δ*CDR20291_1130* deletion does not further the UV defect from the *C. difficile* Δ*sspA* strain alone. Unexpectedly, whenever the double deletion *C. difficile* Δ*sspA* Δ*CDR20291_3080* or the triple deletion *C. difficile* Δ*sspA* Δ*CDR20291_1130* Δ*CDR20291_3080* were tested, they resulted in 10% viability. We hypothesized that these deletions would result in either the same defect of 0.1% viability seen with *C. difficile* Δ*sspA* alone or that UV sensitivity would be increased upon more deletions. This increase in survival would suggest that another factor may be compensating for the loss of these SASPs. Another possibility is that the loss of certain SASPs influences the DNA structure so that UV light affects the strains differently than when those SASPs are present. However, further work is necessary to determine the cause of the partial UV resistance with *C. difficile* Δ*sspA* and Δ*CDR20291_3080* co-deletions. We also found that the other 3 *C. difficile* SASPs, *sspB, CDR20291_1130,* and *CDR20291_3080* are unable to complement a *C. difficile* Δ*sspA* deletion. We hypothesized that this could possibly be due to differences in expression levels. Indeed, when the *sspA* promoter region was used to drive expression of the other SASPs, complementation occurred to approximately 5% of wildtype level. This suggests that the *sspA* promoter region was able to increase expression levels of these SASPs which in turn led to some protection from UV light in comparison to *C. difficile* Δ*sspA* alone. We hypothesize that these other SASPs can bind DNA but need higher expression to do so.

Other analyses revealed that the SASPs SspA, CDR20291_1130, and CDR20291_3080 only minorly contribute to outgrowth. A triple mutation was needed before a ∼2 hour defect was observed. We predict that deletion of all three SASPs depletes the amino acid pool and results in a delay in protein production and outgrowth of the vegetative cell.

In other organisms, SASPs play large roles in protection from chemicals. Interestingly, *C. difficile* SASPs were not important for survival of the spores upon chemical exposure. Nitrous acid was the only chemical where the SASPs may play a minor role in protection. At 5 minutes of exposure, *C. difficile* Δ*sspA* has approximately 3% of wild type survival and this is complemented to wildtype levels. At 10 minutes of exposure, *C. difficile* Δ*sspA* has 8% of wild type survival that, again, complements to wildtype levels. Also at 10 minutes, *C. difficile* Δ*CDR20291_1130* survival is 10% of wildtype and *C. difficile* Δ*CDR20291_3080* is 16%; both could be complemented to wildtype levels. Of note, there were some reductions in spore survival of some mutants compared to wildtype across the other chemicals tested. However, in these cases complementation plasmids did not restore viability, leading to the conclusion that some other factor, besides the SASP deletion, was causing the reduced viability. For evaluating the effect of hypochlorite and glutaraldehyde, reproducibility was an issue. Every trial varied in response to these chemicals, making it difficult to draw a confident conclusion from these data. We conclude that *C. difficile* SASPs do not contribute to chemical resistance. This could be due to protection being evolutionarily unnecessary because *C. difficile* would not typically encounter many of these chemicals in nature or that the spore coat proteins are better suited to resist these conditions. Indeed, the *C. difficile* spore coat is established to contain proteins involved in some resistances [69, 70].

In *B. subtilis*, the Δ*sspA* Δ*sspB* strain is commonly used to characterize the SASPs. It was very surprising that the *C. difficile* Δ*sspB sspA*_G52V_ allele (*sspB*_SNP_) and the *C. difficile* Δ*sspA* Δ*sspB* double mutant were unable to form phase bright spores. Based on the phase-gray cells, we hypothesized that the forespore did not have DPA packaged. Indeed, DPA content is significantly reduced in these SASP-mutant strains. Because DPA contributes to heat resistance and heat resistance is a classically tested feature of endospores and used in sporulation assays, this could have impacted the results of the sporulation assay by killing fully formed spores that do not package DPA and thus exacerbating the sporulation rate phenotype [11, 58, 71–73].

In the *C. difficile* Δ*sspB_SNP_* strain, complementation of the spore phenotype only occurred when *sspB* was driven by a σ^G^ promoter, indicating the importance of expression at the correct time and in the correct compartment during sporulation. The use of the *bcla2* promoter region for σ^E^ was based off Fimlaid *et al.* However, Saujet *et al.* classified *bcla2* as σ^K^ dependent [16, 17]. Either way, mother cell specific promoters do not complement the phenotype, further driving the conclusion that expression must occur in the forespore. This strain could also be complemented by expression of *C. difficile sspA*, suggesting that SspA and SspB may work together.

Due to the phenotype, it was very difficult to purify spores from *C. difficile* Δ*sspB_SNP_* and *C. difficile* Δ*sspA* Δ*sspB*. The spores were treated with lysozyme to digest the mother cell peptidoglycan and release the phase gray spore. These spores are then less dense, further complicating the purification process. Because of these difficulties, the spore amounts used in DPA content assays and immunoblots are rough estimates based on microscope counts. The Coomassie stained gel shows that more protein was present in the *C. difficile* Δ*sspA* Δ*sspB* sample than in the wildtype and the complemented samples even though the immunoblots show similar levels of protein present. This discrepancy could be due to vegetative cells / debris still present after purification through a density gradient or it is possible we underestimated the number of spores in the microscope counts.

Furthermore, evaluation of UV viability on the *C. difficile* Δ*sspB_SNP_* strain complemented with *sspB* shows that these spores have approximately 60% of wildtype viability, a large difference between 0.1% viability in a *C. difficile* Δ*sspA* or the 10% viability of the *C. difficile* Δ*sspB*. Thus, the *sspA*_G52V_ allele does not impair the ability of SspA to protect against UV damage. Due to the observed phenotypes, we hypothesize that SspA and SspB together regulate spore formation. Interestingly, the *C. difficile* Δ*sspA* Δ*sspB* phenotype is similar to the phenotype of a *C. difficile* Δ*spoVT* mutant [17, 74]. In *B. subtilis*, SpoVT is involved in a feed forward loop to regulate sporulation. While it’s expression is through σ^G^, it works to enhance some, and repress other, σ^G^ controlled genes [75]. SpoVT regulates the SASPs, SspA and SspB, with a mutant having a 30% reduction in these proteins [75]. This mutant strain is still able to form phase bright spores but they have a reduced sporulation rate [75]. In *C. difficile*, the *spoVT* mutant has a different phenotype than that observed in *B. subtilis*. This protein is under σ^G^ regulation and possibly also σ^F^ [16, 17, 74]. The *C. difficile* Δ*spoVT* strain forms immature spores that are phase dark [17]. Similarly to *B. subtilis*, SASP expression was reduced in *C. difficile*; *C. difficile sspA* had a 70 fold reduction and *sspB* a 12 fold reduction [17]. The authors suggested that this SASP reduction was probably not sufficient to explain the phenotype [17]. However, our data suggests that the *C. difficile* Δ*spoVT* phenotype may be due to reduced *sspA* and *sspB* levels. This further highlights the differences in sporulation and SASP function in *C. difficile* compared to the model *B. subtilis* and also furthers our hypothesis of SspA and SspB regulating spore formation. Furthermore, in the *B. subtilis* ΔsspA ΔsspB double mutant, expression levels of some genes were changed [76]. However, the expression levels suggest that the SASPs work to negatively regulate multiple forespore genes and even one mother cell gene [76]. They do note that the changes in transcription may not be due to regulation but to actual binding of the SASPs to the genome, which represses transcription. The novel finding of immature spore formation in *C. difficile* Δ*sspA* Δ*sspB* double mutant suggests that *C. difficile* SASPs perform a previously unreported function in the sporulation process. This insight will open new doors for understanding the regulation of sporulation of *C. difficile*.

## Materials and Methods

### Bacterial growth conditions

*C. difficile* strains were grown in a Coy anaerobic chamber at >4% H_2_, 5% CO_2_, and balanced N_2_ at 37 °C in either brain heart infusion supplemented with 5 g / L yeast extract (BHIS) and 0.1% L-cysteine or tryptone yeast medium with 0.1% thioglycolate [77]. When necessary, media was supplemented with thiamphenicol (10 µg / mL), taurocholate (TA) (0.1%), anhydrous tetracycline (100 ng / mL), kanamycin (50 µg / mL), or xylose (1%). *E. coli* strains were grown on LB at 37 °C and supplemented with chloramphenicol (20 µg / mL) for plasmid maintenance. *B. subtilis* BS49 was grown on LB agar or in BHIS broth at 37 °C and supplemented with 2.5 µg / mL chloramphenicol for plasmid maintenance and 5 µg / mL tetracycline during conjugations.

### Plasmid construction

The *C. difficile sspA* targeted CRISPR plasmid was constructed by amplifying 500 bp upstream and downstream from the *sspA* gene using primer pairs 5’sspA_MTL, 3’sspA_up and 5’sspA_down, 3’sspA_downMTL, respectively. The fragments were cloned into the *Not*I and *Xho*I site of pKM126 by Gibson assembly [78]. The gRNA was retargeted to *sspA* by inserting gBlock CRISPR_sspA_165 into the *Kpn*I and *Mlu*I sites by Gibson assembly, generating the pHN05 plasmid. The *spl, sspB, CDR20291_1130* and *CDR20291_3080* targeting plasmids were similarly constructed. For *spl* plasmid construction, 5’ spl_UP and 3’ spl_UP amplified the upstream homology arms and the downstream homology by 5’ spl_DN and 3’ spl_DN. These were cloned into pKM197 at the *Not*I and *Xho*I sites. The gRNA was retargeted by cloning into the *Kpn*I and *Mlu*I sites gBlock CRISPR_spl_647 resulting in plasmid pHN61. Primers 5’ sspB UP and 3’ sspB UP amplified the upstream region, while primers 5’ sspB DN and 3’ sspB DN amplified the downstream portion. These fragments were also cloned into pKM126 at the same sites as previously used for homology. The gRNA was switched with CRISPR_sspB_144 at *Kpn*I and *Mlu*I sites, generating pGC05. For *CDR20291_1130,* primer pairs 5’CDR20291_1130_UP, 3’ CDR20291_1130_UP and 5’CDR20291_1130_DN, 3’CDR2021_1130_DN were used, respectively, to amplify upstream and downstream portions of *CDR20291_1130* homology. For *CDR20291_3080,* primer pairs 5’CDR20291_3080_UP, 3’ CDR20291_3080_UP and 5’CDR20291_3080_DN, 3’CDR2021_3080_DN were used, respectively, to amplify upstream and downstream portions of *CDR20291_3080* homology. These homology arms were cloned by Gibson assembly into pKM126 [52] at sites *Not*I and *Xho*I [78]. gRNAs CRISPR_CDR20291_1130_114 and CRISPR_CDR20291_3080_184 were cloned into the plasmids at the *Kpn*I and *Mlu*I sites. Next, because of issues with the *tet* promoter (in pKM126) causing leaky expression of *cas9*, and causing potential problems during conjugation, *tetR* was replaced with *xylR* from pIA33 [79]. The *xylR* region was amplified with primers 5’ sspB.xylR and 3’cas9_Pxyl2 for sspB plasmid, 5’CD1130_HR_xylR and 3’cas9_Pxyl2 for the CDR20291_1130 plasmid and the primers 5’CD3080_HR_xylR and 3’cas9_Pxyl2 for the CDR20291_3080 plasmid to create pHN101, targeting *sspB*, pHN32, targeting *CDR20291_1130,* and pHN34, targeting *CDR20291_3080*.

For *C. difficile* Δ*sspA* complementation plasmids, the genes and promoter regions were inserted by Gibson assembly into the *Not*I and *Xho*I sites of pJS116 [53] by Gibson assembly [78]. The *sspA* gene plus 500 bp upstream was amplified using primer pair 5’sspA_MTL and 3’ sspA.pJS116 to create plasmid pHN11. The vector, pHN14, consisting of the *sspB* gene and 500 bp upstream was generated using primers 5’ sspB UP and 3’sspBpJS116. The *sspA* portion of the *sspA* and *sspB* double mutant complementation vector, pHN30, was amplified with 5’sspApJS116 and 3’ sspAsspB. The *sspB* portion of the double mutant complement was amplified with 5’ sspAsspB and 3’ sspBpJS116. For pHN84, the *sspA* complement with a 6x histidine tag on the C-terminus, the primers 5’sspA_MTL and 3’ sspA.His_pJS116 were used. The primers 5’ sspB UP with 3’ PsspB_sspA and 5’ PsspB_sspA with 3’ sspA.pJS116 were used to generate a plasmid with 500 bp upstream of the *sspB* gene driving expression of the *sspA* gene for pHN91. The 500 bp upstream of *sspA* was amplified with 5’sspA_MTL and 3’sspB_sspA and used to drive expression of the *sspB* gene, amplified with 5’sspB_sspA and 3’sspBpJS116 for the pHN83 complementation vector. To generate the pHN56 vector of the *CDR20291_1130* gene and 500 bp upstream, the primer pair 5’ 1130comp and 3’ 1130comp were used. For amplification of the *CDR20291_3080* gene and 500 bp upstream, pHN57, the primers 5’ 3080comp and 3’ 3080comp were used. Generation of 500 bp upstream of *sspA* was amplified with primers 5’sspA_MTL and 3’ sspA_CD1130 and used to drive expression of the *CDR20291_1130* gene amplified with primers 5’ CD1130_sspA and 3’ 1130comp for plasmid pHN96. The plasmid pHN97 was generated with 500 bp upstream of *sspA*, with primers 5’sspA_MTL and 3’ sspA_CD3080 and the *CDR20291_3080* gene amplified with 5’ CD3080_sspA and 3’ 3080comp. For the negative control, pHN109, of *mCherry* driven by *sspA* promoter region, the *sspA* promoter region was amplified with primers 5’sspA_MTL and 3’ PsspA.mCherry, while the *mCherry* gene was amplified from pRAN473 [80] with primers 5’ PsspA.mCherry and 3’ mCherry.PsspA. Another negative control of just the sspA promoter region, pHN102, was generated by using 5’sspA_MTL and 3’ PsspA.pJS116 to amplify the *sspA* promoter region. To generate the σ^E^_sspB plasmid, the promoter region of *bclA*2 was amplified with 5’ sigE.bclA2_pJS116 and 3’ sigE.bclA2_sspB and the *sspB* gene with 5’ sigE.bclA2_sspB and 3’ sspB_pJS116, creating pHN80. For the σ^K^_sspB complement vector, the promoter region of *sleC* was amplified with 5’ pJS116_sigK and 3’ sigK_sspB and the *sspB* gene with 5’ sigK_sspB and 3’sspB_pJS116, generating pHN49. The promoter region of *gpr* was amplified with 5’ pJS116_sigF and 3’ sigF_sspB and the sspB gene with 5’ sigF_sspB and 3’ sspB_pJS116 to generate the σ^F^_sspB plasmid, pHN47. All plasmid sequences were confirmed by DNA sequencing. Cloning was done in *E. coli* DH5α. A complete list of oligonucelotides used and the strains and plasmids generated can be found in Tables S1 and S2, respectively.

### Conjugations

The resulting plasmids, pGC05, pHN05, pHN61, pHN101, pHN32, and pHN34 were conjugated separately into *C. difficile* R20291 using *B. subtilis* BS49 as a conjugal donor, as described previously [52]. Briefly, 0.25 mL of *C. difficile* R20291 overnight culture was back diluted into 4.75 mL fresh BHIS and incubated for 4 hours. Meanwhile, 5 mL of BHIS supplemented with chloramphenicol and tetracycline was inoculated with 1 colony of *B. subtilis* BS49 containing the plasmid and was incubated at 37 °C for 3 hours. After incubation, the *B. subtilis* culture was passed into the anaerobic chamber and 100 µL culture was plated on TY agar medium, along with 100 µL of *C. difficile* R20291. This was incubated for approximately 24 hours. Growth was then suspended in 1.5 mL of BHIS and plated onto BHIS agar supplemented with thiamphenicol and kanamycin for selection. Resulting transconjugant colonies were streaked, twice, onto BHIS supplemented with thiamphenicol and kanamycin (TK) or BHIS supplemented with tetracycline (to screen for the conjugal transfer of the *Tn*916 transposon). Thiamphenicol resistant and tetracycline sensitive colonies were tested by PCR for plasmid components.

### CRISPR induction

Overnight cultures of 5 mL TY medium supplemented with thiamphenicol were inoculated with one colony of a *C. difficile* R20291 strain containing a CRISPR mutagenesis plasmid with *tet*-driven *cas9* (pGC05 and pHN05). After approximately 16 hours of growth, 0.25 mL of overnight culture was back diluted with 4.75 mL TY broth and supplemented with thiamphenicol and anhydrous tetracycline and incubated for 6 hours [52]. A loopful (approximately 10 µL) of culture was plated onto BHIS and the individual colonies were tested by PCR for the mutation. Once PCR confirmed, the plasmid was cured by passaging onto BHIS plates. For induction of *xyl*-driven *cas9* plasmids (pHN61, pHN101, pHN32 and pHN34), colonies were passaged 3 times on TY agar supplemented with 1% xylose and thiamphenicol. Mutants were detected by PCR and the plasmid was cured by passaging in BHIS broth supplemented with 0.5% xylose. Pairwise mutants were generated by conjugating the appropriate plasmid into the necessary mutant strain and inducing as described above.

### Spore Purification

Cultures were plated onto 70:30 media or CDMM minimal media where indicated and incubated 5 days. Spores were purified as previously described [53, 57, 58, 81]. Briefly, plates were scraped and contents suspended in dH_2_O overnight at 4 °C. The pellets were resuspended then centrifuged at max speed. The upper, fluffy-white, layer was removed and resuspended again in dH_2_O. This process was repeated approximately 5 times. The spores were then separated by density gradient in 50% sucrose solution at 3,500 xg for 20 minutes. The spore pellet was washed approximately 5 times in dH_2_O. The spores were stored at 4 °C until use.

To purify the Δ*sspB_SNP_* and the Δ*sspA* Δ*sspB* double mutant spores, the pellets were scraped into dH_2_O and left overnight at 4 °C. The pellets were then resuspended with 1 µg of lysozyme and incubated for 4 hours at 37 °C. The suspension was centrifuged at max speed for 1 minute and the upper phase removed, and the pellet resuspended with dH_2_O. This process was repeated approximately 5 times before 5 mL of spores were layered onto a HistoDenz gradient of 10 mL of 50% and 10 mL of 25%. This was centrifuged 30 minutes at 18,900 xg at 4 °C. The pellet was then washed approximately 5 times in dH_2_O and stored at 4°C until use.

### Germination assay and DPA content

Germination was monitored using a Spectramax M3 plate reader (Molecular Devices, Sunnyvale, CA). 5 µL of OD_600_ = 100 spores were added to 95 µl germination buffer consisting of a final concentration 1x HEPES, 30 mM glycine, 10 mM TA and the OD_600_ was monitored for 1 hour at 37 °C. To assay total DPA content, 1 x 10^6^ spores in 20 µL, were boiled at 95 °C for 20 minutes. 5 µL (an equivalent of 2.5 x 10^5^ spores) of the solution was added to 95 µL of 1X HEPES buffer with 250 µM TbCl_3_ and analyzed by excitation at 275 nm and emission at 545 nm with a 420 nm cutoff [56, 57, 61, 72, 82].

### Western blotting

For the Δ*sspB_SNP_* strain, approximately 1 x 10^7^ spores were incubated for 1 hour in 50 µL of BHIS with or without 10 mM TA at 37 °C. The solutions were boiled for 20 minutes in 2x NuPAGE buffer at 95 °C. 10 µL (equivalent to approximately 1 x 10^6^ spores) of each solution was separated on a 15% SDS PAGE gel. For the double mutant strain, Δ*sspA* Δ*sspB*, approximately 4-8 x 10^4^ total spores were separated. The protein was transferred to polyvinylidene difluoride (PVDF) membranes at 0.75 amps for 1.5 hours for the Δ*sspB_SNP_* strain and at 1 amp for 30 minutes for the Δ*sspA* Δ*sspB* strain. The membranes were blocked overnight in 5% milk in TBST. The following day, the membranes were washed 3 times, 15 minutes each, at room temperature. SleC anti-sera was added to 5% milk in TBST at a 1:20,000 dilution for 1 hour. The membranes were washed again, as above. The anti-rabbit secondary antibody was diluted to 1:20,000 in 5% milk in TBST and incubated for 1 hour. The membranes were again washed, as above. The membranes were incubated with Pierce ECL Western Blotting Substrate (ThermoScientific) for 1 minute and then x-ray film was exposed and developed.

### Phase contrast imaging

Strains were plated onto 70:30 (70% SMC medium and 30% BHIS medium) sporulation media and incubated for 6 days. After the incubation period, the growth was harvested and suspended in fixative (4% formaldehyde, 2% glutaraldehyde in 1x PBS). The samples were imaged on a Leica DM6B microscope at the Texas A&M University Microscopy Imaging Center.

### Sporulation

Sporulation assays were completed as previously described [73]. Briefly, three 70:30 plates were inoculated with fresh colonies of the indicated strains. These were grown for 48 hours before half of the plate was harvested into 600 µL of pre-reduced PBS solution and the pellets resuspended. 300 µL of suspension was removed from the chamber and heated for 30 minutes at 65 °C and mixed every 10 minutes by inversion. After heating, these samples were passed into the anaerobic chamber for enumeration. Both the heat-treated and the remaining 300 µL of untreated suspension was serially diluted and plated in technical triplicates of 4.5 µL spots on rich, BHIS, medium supplemented with 0.1% TA. After 22 hours of growth, the CFUs were enumerated. The ratios of treated to untreated were calculated and then the efficiency of sporulation determined in comparison to the wildtype strain. The experiment was performed in biological triplicate.

### UV exposure

The 302 nm UV lamp (Fisher Scientific) was allowed to warm-up for 20 minutes before UV exposure. Spores were diluted in PBS to 1 x 10^7^ spores / mL. A sample was collected at T_0_ for initial, untreated, spore calculations. 1 mL of a spore solution was added to a miniature glass (UV penetrable) petri dish. These dishes were placed under the UV lamp and exposed with constant agitation for the indicated time. After exposure, the untreated and treated samples were serially diluted then introduced into the anaerobic chamber where they were plated onto BHIS medium supplemented with taurocholic acid and thiamphenicol, for plasmid maintenance. After 48 hours of incubation, the CFUs were enumerated. Treated spore counts were normalized to untreated and then this ratio was normalized to the ratio for wildtype spores.

### Outgrowth

Outgrowth, post-germination, was performed in half-strength BHIS supplemented with 30 mM glycine and 10 mM taurocholic acid. Thiamphenicol was supplemented where necessary to maintain plasmids. Spores were added to an OD_600_ of 0.05 and the OD_600_ was recorded over time.

### Chemical resistance

1 x 10^7^ spores were suspended in PBS with the indicated chemical concentration for various exposure times. The exposed spores were immediately serially diluted into PBS tubes and plated onto rich media supplemented with taurocholate. For generation of nitrous acid, a one to one volume of 1 M sodium acetate, pH 4.5 and 1 M sodium nitrite were combined and incubated for at least 30 minutes before spore exposure. For formaldehyde exposure, the exposed solution was serially diluted into PBS with 400 mM glycine and incubated for at least 20 minutes to quench the formaldehyde before plating.

## Acknowledgments

We thank Ge Chen for the construction of the pGC05 plasmid and members of the Sorg laboratory for critical comments during preparation of this manuscript. This project was supported by awards R01AI116895 and U01AI124290 from the National Institute of Allergy and Infectious Diseases. The content is solely the responsibility of the authors and does not necessarily represent the official views of the NIAID. The funders had no role in study design, data collection and interpretation, or the decision to submit the work for publication.

---

**Supplement Figure 1.**
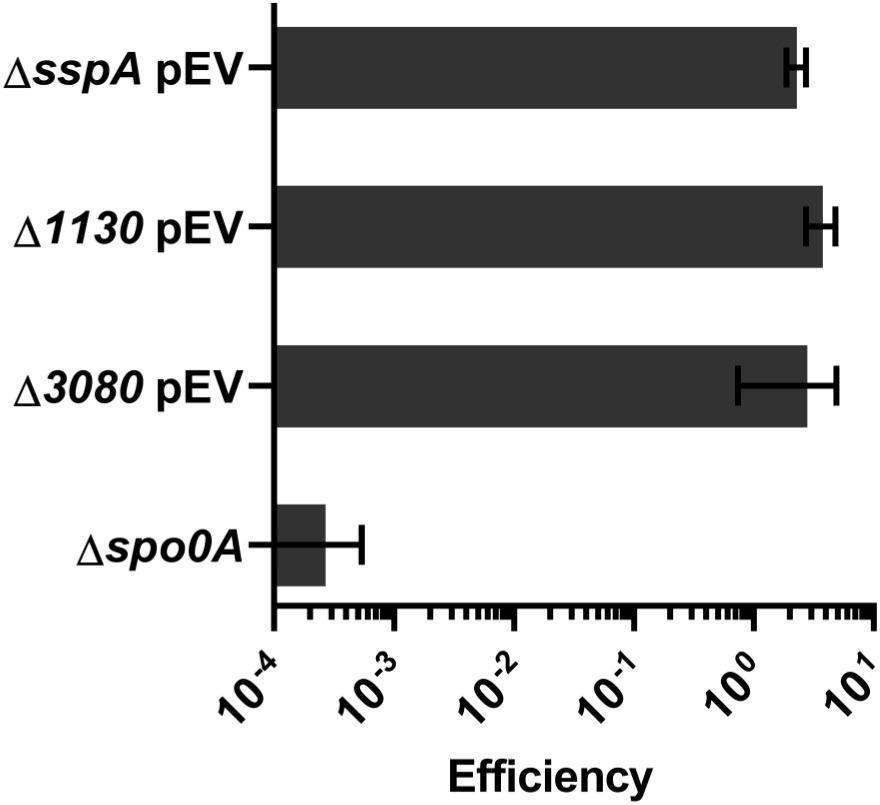
Sporulation efficiency is not affected by deletion of *sspA, CDR20291_1130* or *CDR20291_3080* alone. Strains were grown on sporulation medium for two days. Sporulating cultures were heat treated at 65 °C. Sporulation rate was determined by comparison of the CFU of heat treated culture to CFU of untreated culture and then the ratios were compared to wildtype. pEV indicates an empty vector. All data represents the average of three independent experiments. Statistical analysis by one way ANOVA with Dunnett’s multiple comparison test.

**Supplement Figure 2.**
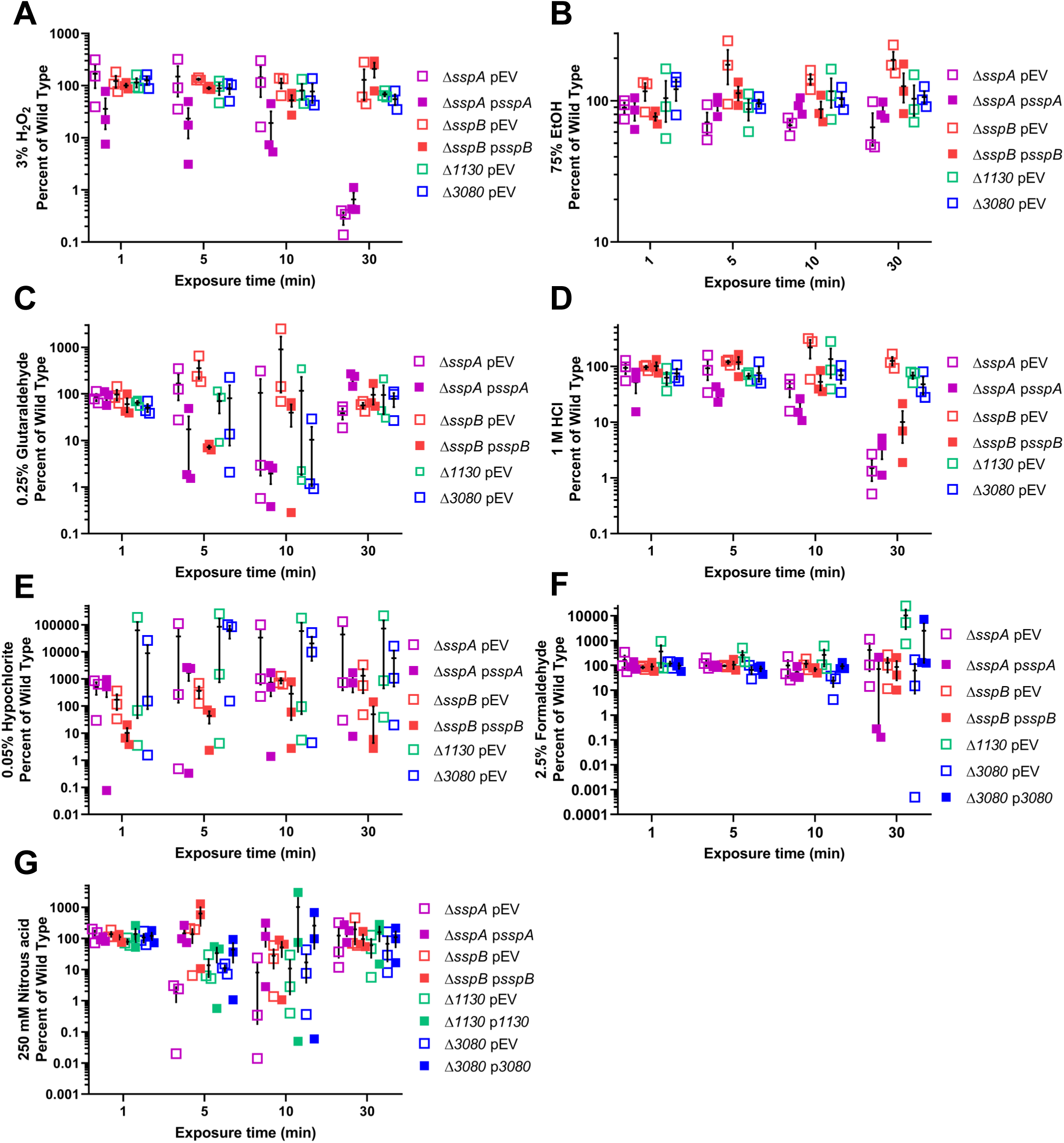
Chemical resistance is not impacted by loss of individual SASPs. 1 x 10^7^ spores were exposed to chemicals for 1, 5, 10, or 30 minutes. After exposure, solutions were serially diluted and plated onto rich medium with germinants. The CFUs were enumerated and compared to unexposed samples and then this ratio was compared with that of the wildtype spores. A) 3% H202 B) 75% EtOH C) 0.25% Glutaraldehyde D) 1 M HCL E) 0.05% hypochlorite F) 2.5% Formaldehyde G) 250 mM Nitrous Acid. pEV indicates an empty vector. All data represents the average of three independent experiments. Statistical analysis by two way ANOVA with Dunnett’s multiple comparison. A) *C. difficile* Δ*sspA* pEV and *C. difficile* Δ*sspA* p*sspA* P<0.0001 at 30 minutes. C) P<0.001 for *C. difficile* Δ*sspB* p*sspB* 5 minutes and *C. difficile* Δ*sspA* p*sspA* at 10 minutes. P<0.05 for *C. difficile* Δ*CDR20291_3080* pEV at 10 minutes. D) P<0.05 for *C. difficile* Δ*sspA* p*sspA* at 5 minutes and *C. difficile* Δ*sspB* p*sspB* at 30 minutes. P<0.01 for *C. difficile* Δ*sspA* p*sspA* at 10 minutes. P<0.001 for *C. difficile* Δ*sspA* pEV and *C. difficile* Δ*sspA* p*sspA* at 30 minutes. E) P<0.01 for *C. difficile* Δ*sspB* p*sspB* at 1 minute. F) P<0.05 for *C. difficile* Δ*CDR20291_3080* at 10 minutes. G) P<0.05 for *C. difficile* Δ*CDR20291_1130* at 5 minutes, *C. difficile* Δ*sspA* pEV and *C. difficile* Δ*CDR20291_1130* pEV at 10 minutes. P<0.01 for *C. difficile* Δ*CDR20291_3080* at 5 minutes. P<0.001 for *C. difficile* Δ*sspA* pEV at 5 minutes.

**Supplement Figure 3.**
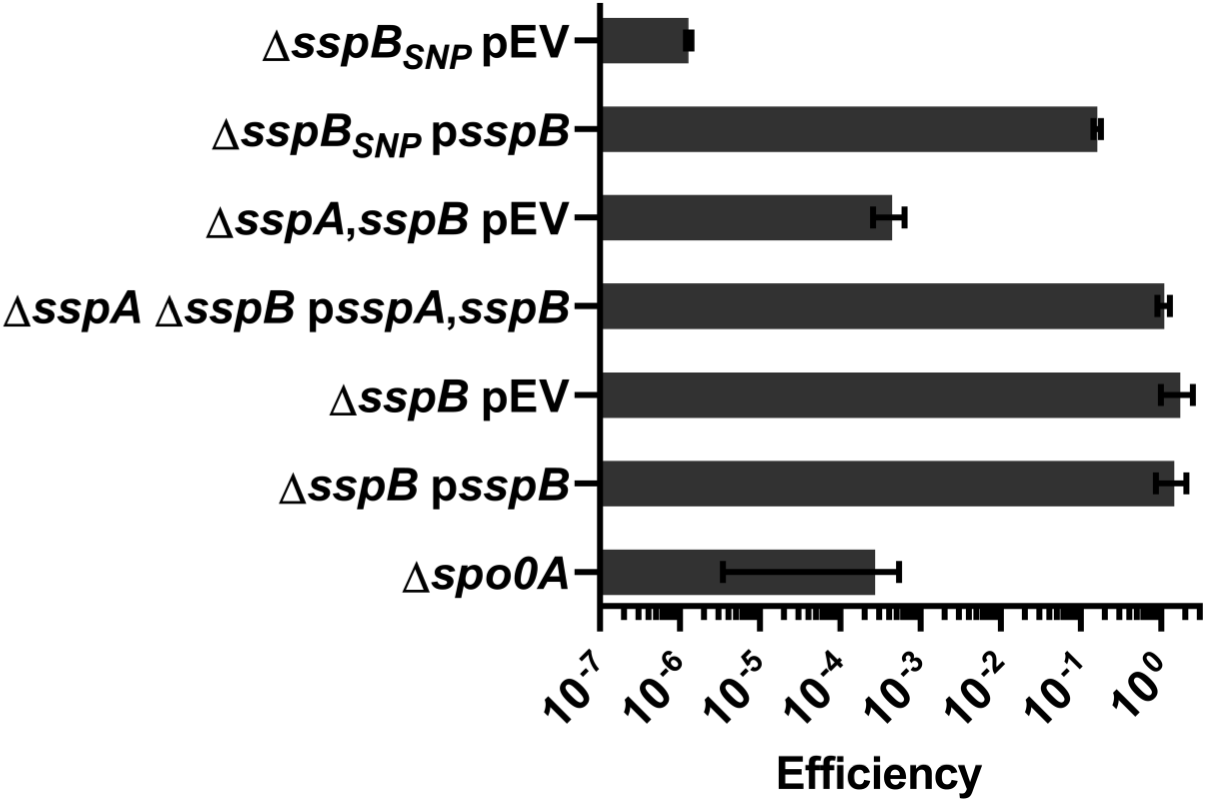
Sporulation efficiency of mutant strains. Strains were grown on sporulation medium for two days. Sporulating cultures were heat treated at 65 °C. Sporulation rate was determined by taking the ratio of the CFU of heat treated culture to CFU of untreated culture and then the ratios were compared to wildtype. pEV indicates an empty vector. All data represents the average of three independent experiments. Statistical analysis by one way ANOVA with Dunnett’s multiple comparison test.

**Supplement Figure 4.**
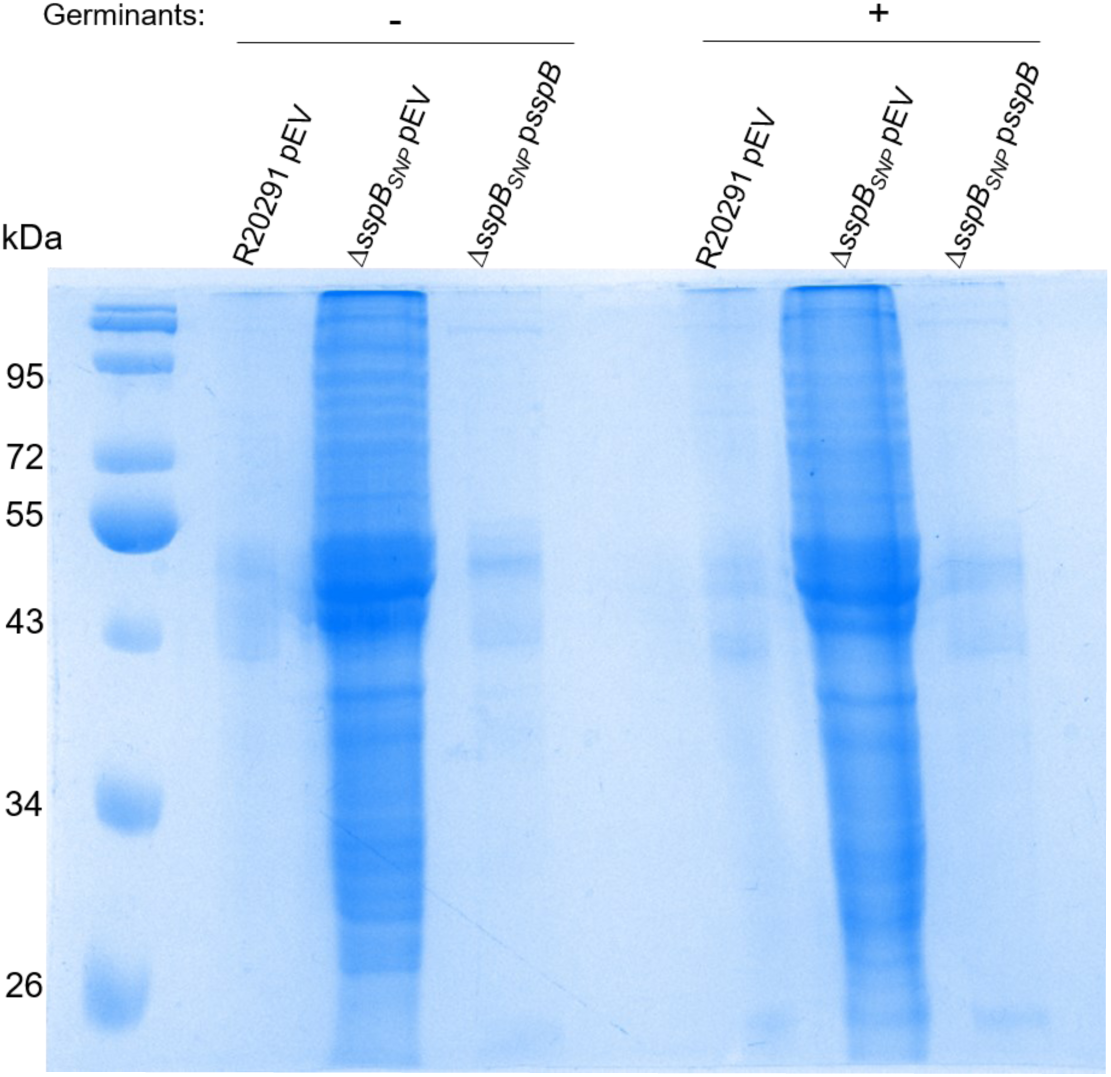
Coomassie-stained gel of Δ*sspA* Δ*sspB* samples. The same sample volumes used for the Δ*sspA* Δ*sspB* SleC cleavage assay were separated by a 15% SDS PAGE and stained with Coomassie. pEV indicates an empty vector.

**Supplement Table 1.**
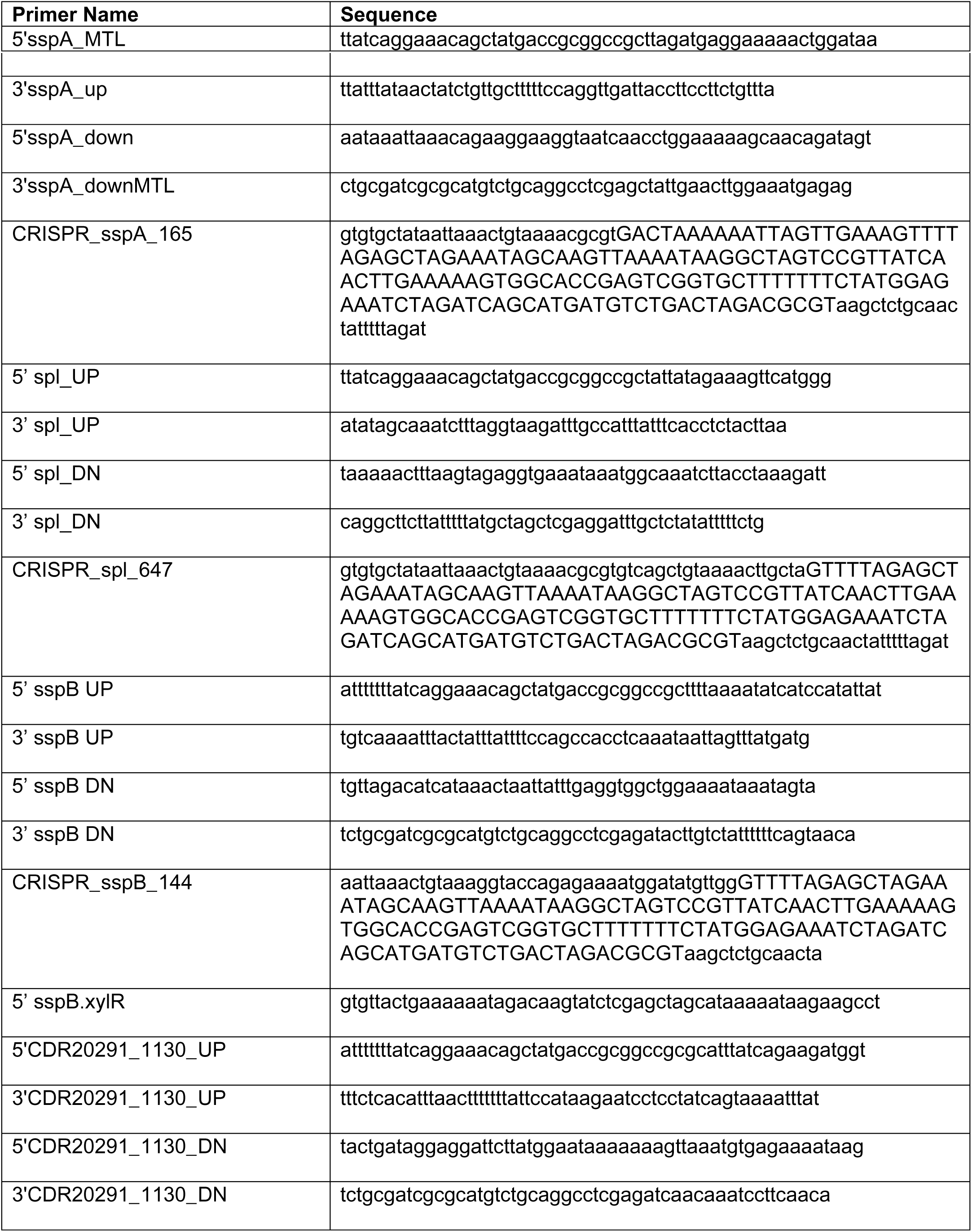
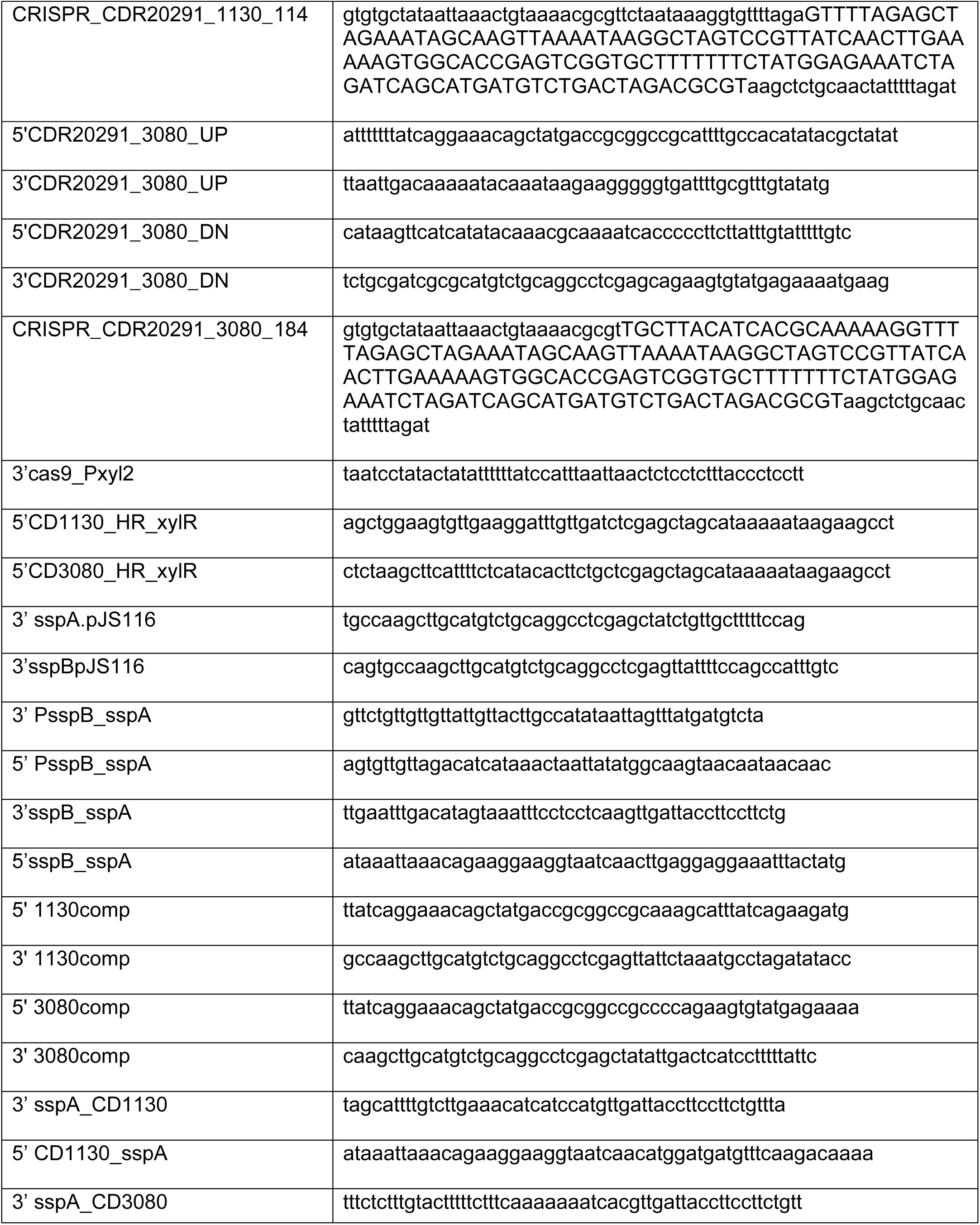
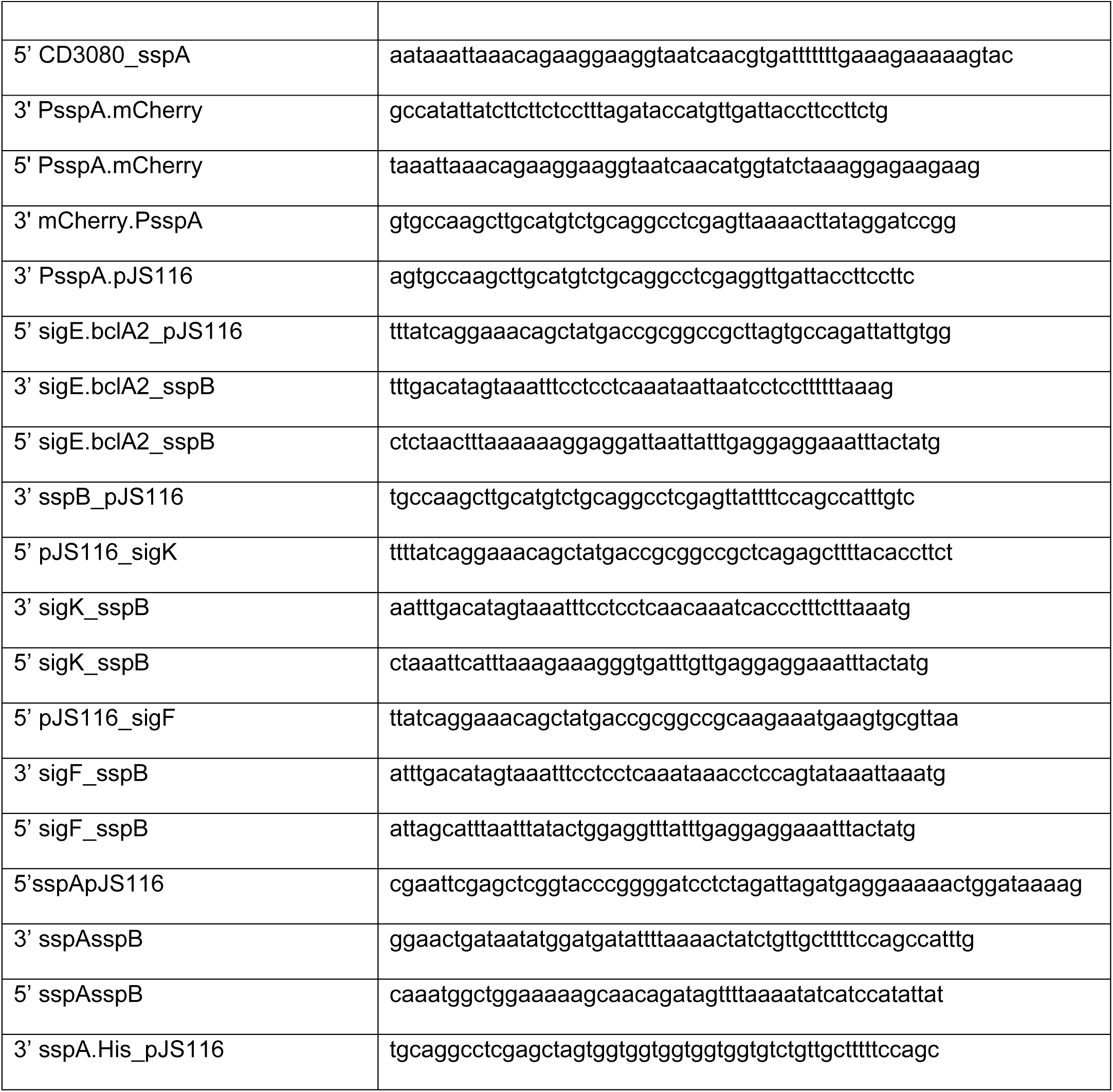
Primers used in this study.

**Supplement Table 2.**
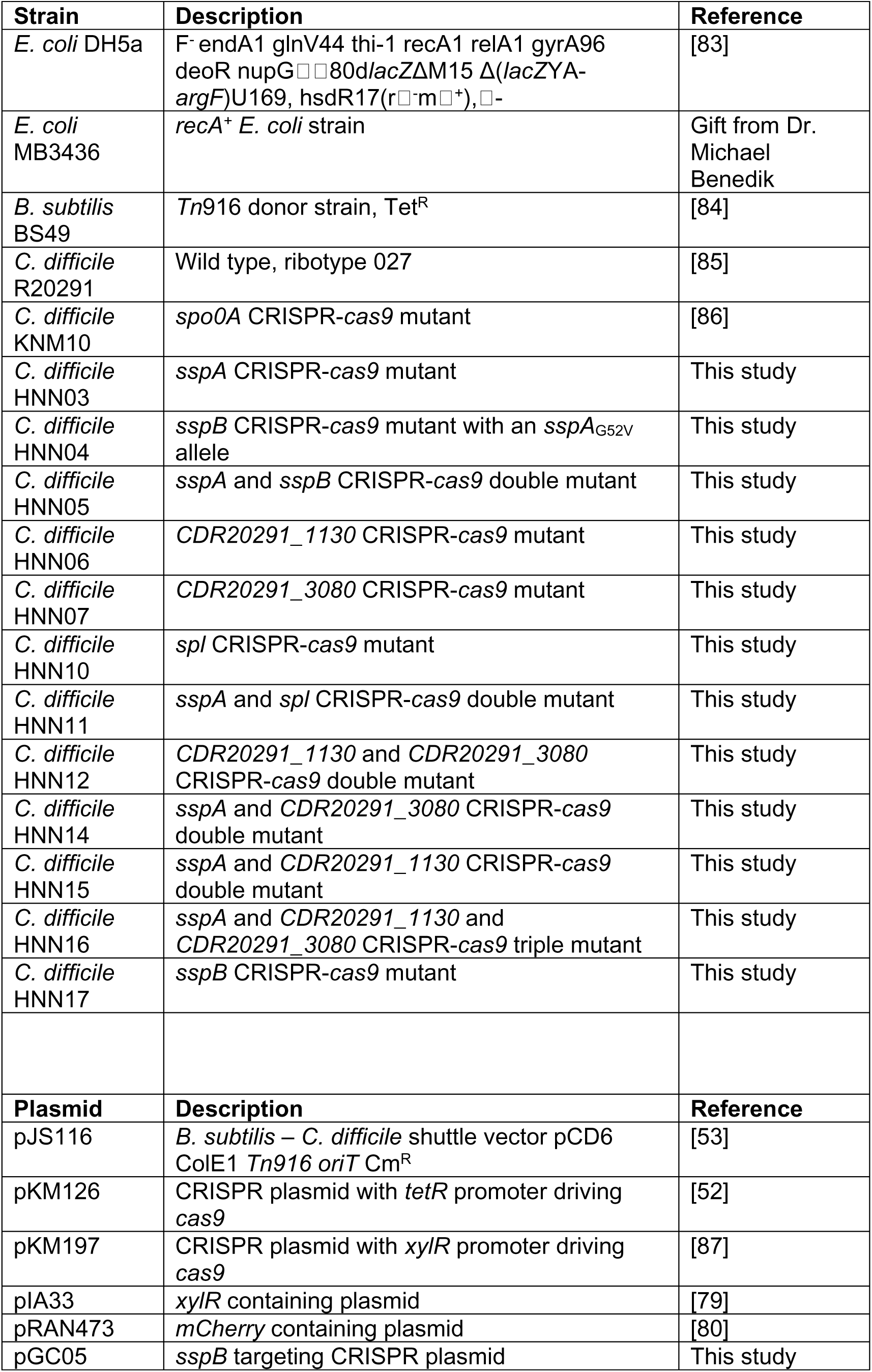
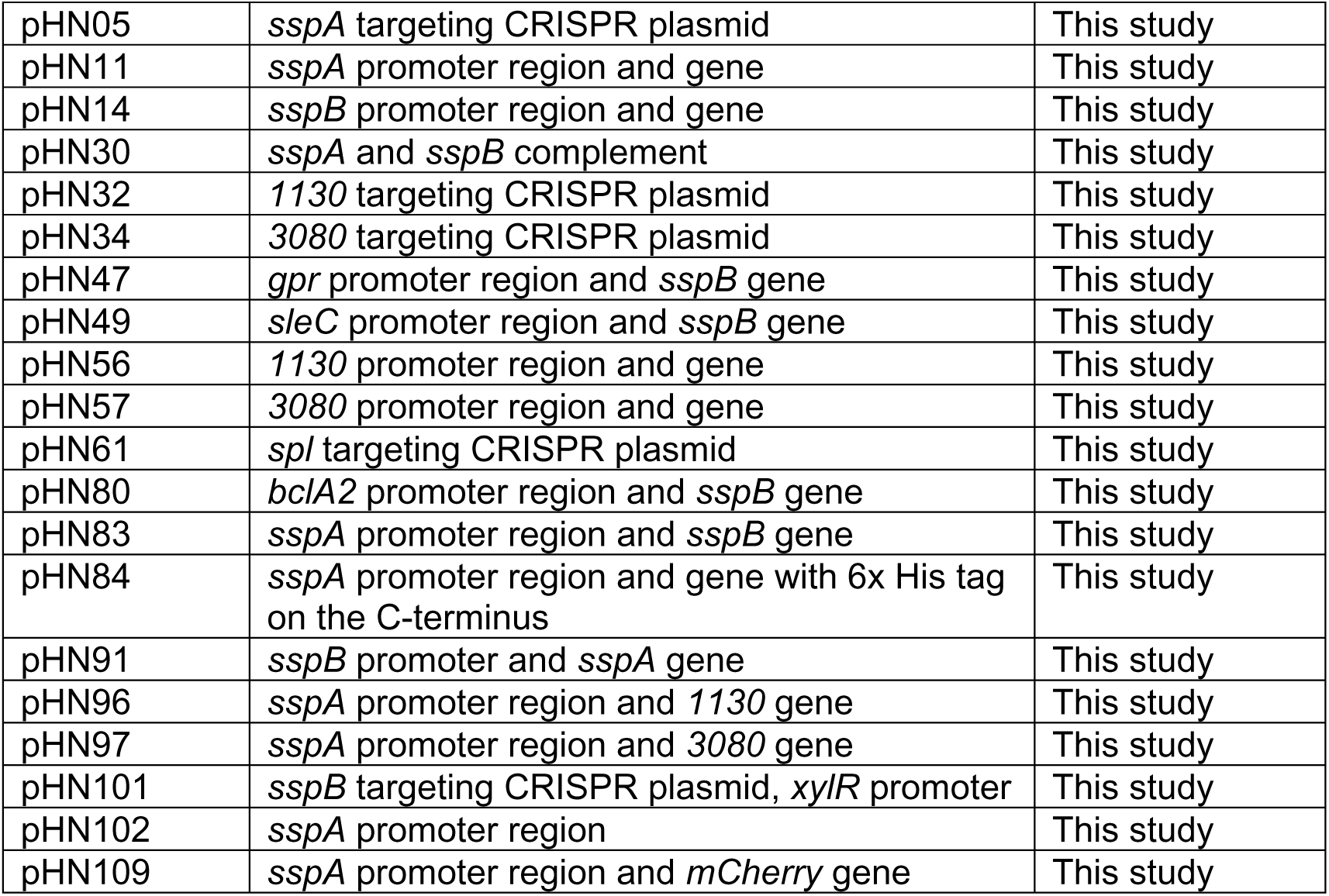
Strains and plasmids used in this study.

## Notes

### Competing Interest Statement

The authors have declared no competing interest.

